# An ecological basis for dual genetic code expansion in marine deltaproteobacteria

**DOI:** 10.1101/2021.03.15.435355

**Authors:** Veronika Kivenson, Blair G. Paul, David L. Valentine

**Author notes:** **Correspondence:** David L. Valentine. VK: Oregon State University, Corvallis, OR 97331. BGP: Marine Biological Laboratory, Woods Hole, MA 02543.

## Abstract

Marine benthic environments may be shaped by anthropogenic and other localized events, leading to changes in microbial community composition evident decades after a disturbance. Marine sediments in particular harbor exceptional taxonomic diversity and can shed light on distinctive evolutionary strategies. Genetic code expansion may increase the structural and functional diversity of proteins in cells, by repurposing stop codons to encode noncanonical amino acids: pyrrolysine (Pyl) and selenocysteine (Sec). Here, we show that the genomes of abundant Deltaproteobacteria from the sediments of a deep-ocean chemical waste dump site, have undergone genetic code expansion. Pyl and Sec in these organisms appear to augment trimethylamine (TMA) and one-carbon metabolism, representing key drivers of their ecology. The inferred metabolism of these sulfate-reducing bacteria places them in competition with methylotrophic methanogens for TMA, a contention further supported by earlier isotope tracer studies and reanalysis of metatranscriptomic studies. A survey of genomic data further reveals a broad geographic distribution of a niche group of similarly specialized Deltaproteobacteria including at sulfidic sites in the Atlantic Ocean, Gulf of Mexico, Guayamas Basin, and North Sea, as well as in terrestrial and estuarine environments. These findings reveal an important biogeochemical role for specialized Deltaproteobacteria at the interface of the carbon, nitrogen and sulfur cycles, with their niche adaptation and ecological success seemingly enabled by genetic code expansion.

## Introduction

Marine sediments harbor a diverse and dense microbiome (Sogin et al., 2006; Orcutt et al., 2011) that has evolved over billions of years to perform a complex network of biogeochemical processes key to the carbon, nitrogen and sulfur cycles. These environments now also serve as a repository for industrial, military and municipal waste (Council on Environmental Quality, 1970; Duedall et al., 1983), which are expected to impart localized ecological and evolutionary pressures. One form of adaptation relevant to marine microorganisms is genetic code expansion, a type of code flexibility in which protein synthesis occurs with more than the twenty canonical amino acids. Known variants code for the 21st and 22nd amino acids, selenocysteine (Sec) and pyrrolysine (Pyl), respectively, through the repurposing of stop codons.

In bacteria, Sec is inserted in polypeptides at an in-frame UGA (opal / stop) codon, via recognition of a downstream ∼50 nt hairpin structure known as a Sec insertion sequence (Zinoni et al., 1990; Liu et al., 1998). Proteins containing Sec occur in a subset of organisms in each of the three domains of life, including many bacteria such as members of Proteobacteria, Chloroflexi, and Firmicutes (Böck et al., 1991; Low and Berry, 1996; Böck, 2000). Sec is part of the active site of a subset of redox enzymes involved in energy metabolism, conferring a higher catalytic efficiency (Cone et al., 1976; Zinoni et al., 1987; Mukai et al., 2016); for example, when Sec is replaced by its counterpart, cysteine, in formate dehydrogenase, the rate of oxidation is slowed by approximately three orders of magnitude (Axley et al., 1991). These proteins have critical roles in marine biogeochemical cycles (Copeland, 2005; Zhang and Gladyshev, 2008) and microbial Sec-utilization may be facilitated in aquatic / marine environments because of a sufficient supply of the required element, Selenium, in the ocean (Peng et al., 2016).

Pyl is encoded in place of an in-frame UAG (amber / stop) codon and occurs in <1% of sequenced organisms, primarily methanogenic archaea (Atkins and Gesteland, 2002; Srinivasan et al., 2002a; Gaston et al., 2011a). The Pyl residue is crucial as part of the active site of mono-, di-, and tri-methylamine methyltransferases (Paul et al., 2000; Atkins and Gesteland, 2002; Gaston et al., 2011b). Methylated amines are ubiquitous in marine sediment, the water column, and marine aerosols, and play a significant role in ocean carbon and nitrogen cycles (Sun et al., 2019). Trimethylamine (TMA) is an important compound in this family, with concentrations ranging from a few nM in the water column, to low µM in sediment pore water, representing a significant pool of dissolved organic nitrogen (Chen, 2012; Sun et al., 2019). Along with their precursor quaternary amines, methyl amines contribute significantly to the production of methane (Borges et al., 2016).

Although anaerobic metabolism of methylated amines is commonly attributed exclusively to methanogenic archaea, genetic code expansion with Pyl in bacteria was identified in the TMA-utilizing *Acetohalobium arabaticum* (Prat et al., 2012), in human gut bacteria (Kivenson and Giovannoni, 2020), and in uncultivated marine worm symbionts that also encode Sec (Zhang and Gladyshev, 2007). However, the occurrence, distribution, and metabolic significance of genetic code expansion in bacteria from marine environments remains unknown.

In this study, we capitalize on our recent discovery of a deep ocean industrial waste dump (Kivenson et al., 2019) to investigate the perturbed sediment microbiome and examine genetic code expansion which may enable ecological success associated with this disturbance. In the dominant Deltaproteobacteria, we find extensive evidence of a largely unexplored mechanism for TMA and one-carbon metabolism augmented by dual genetic code expansion. We characterize these Sec-and Pyl-dependent metabolic pathways and identify additional occurrences of these adaptations in widely distributed Deltaproteobacteria from genomic, metagenomic, and transcriptomic data, uncovering an important role for these organisms in coupling the carbon, nitrogen and sulfur cycles in oxygen-limited marine environments.

## Results and Discussion

### An Industrial Waste Microbiome

Photo-surveys using an autonomous underwater vehicle revealed approximately sixty of an estimated half a million barrels dumped in the San Pedro Basin, off the coast of California (**Fig. S1**), as described in our previous study (Kivenson et al., 2019). A subset of the barrels exhibited a ring-form microbial mat that commonly, but not exclusively, encircled the barrels. Burrows were present throughout the seafloor and notably absent between each barrel and its microbial mat ring (**Fig. S2**). A survey of 16S rRNA genes was conducted for the upper 2cm of sediment at the microbial mats, just outside of these mats, and ≥60m from any visible barrels, to determine the identity and relative abundances of microbial taxa associated with these habitats. The circular mat rings were sampled at barrel 16 (bbl-16) (**Fig. S3, Fig. 1A**), and barrel 31 (bbl-31) (**Fig. 1B, 1C)**. These microbial mats were less diverse than surrounding sediments (**Fig. 1D, Fig. S4**), consistent with a response to a major environmental disturbance.

**Figure 1.**
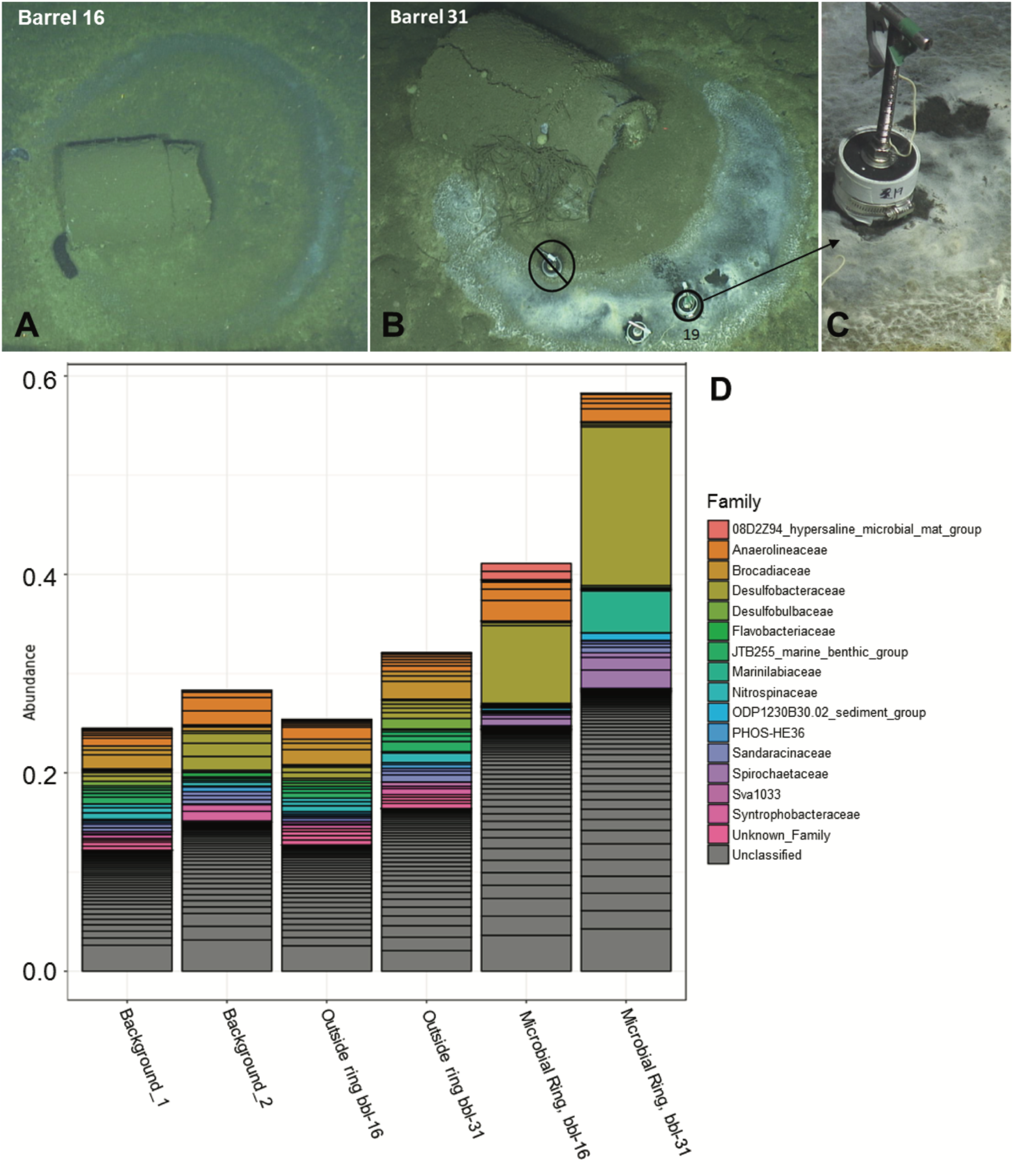
**A)** Barrel 16 encircled by a microbial mat ring **B)** Barrel 31 with microbial mat ring; sediment collection site of Core 19. **C)** Close-up of Core 19 at barrel 31 microbial mat **D)** Abundance of top taxa by sample for background, outside ring, and microbial ring samples.

The most abundant taxa for both of the microbial mat samples (ASV-1) is a member of the *Desulfobacteraceae* family, most closely affiliated with the *Desulfobacula* genus, and comprises 16% and 8% of the total bacterial community for bbl-31 and bbl-16, respectively (**Fig. 1D, Table S1**). These abundances are substantially greater than that of the most abundant organism, a member of the SEEP-SRB1 clade of *Desulfobacteraceae*, present at 3.2% in surface sediment at the nearby San Pedro Ocean Times Series (SPOTS) station (Monteverde et al., 2018). As observed in the SPOTS sediment dataset, sediments (non-ring samples) did not have a single taxa exceeding 4% of the total community (**Table S1**).

### Genome Reconstruction of Enriched Taxa

Metagenomic sequencing was performed for the six sediment samples, with between 9 and 22 GB of raw sequencing reads generated per sample (**Table S2**). In total, eleven genomes were reconstructed from the two microbial mat samples, of which six are estimated to be near-complete (90+%) (**Table 1, Table S3, S4**). The reconstructed genomes include members of the *Desulfobacterales, Bacteroidales, Gemmatimondales, Victivalles*, and *Spirochaetales* orders, as well as several candidate phyla: *Zixibacteria, Woesarchaeota, Marinimicrobia*, and *Latescibacteria*. Many of these taxa are commonly detected in oxygen-limited marine sediment, including the SPOTS data set (Monteverde et al., 2018).

**Table 1.**
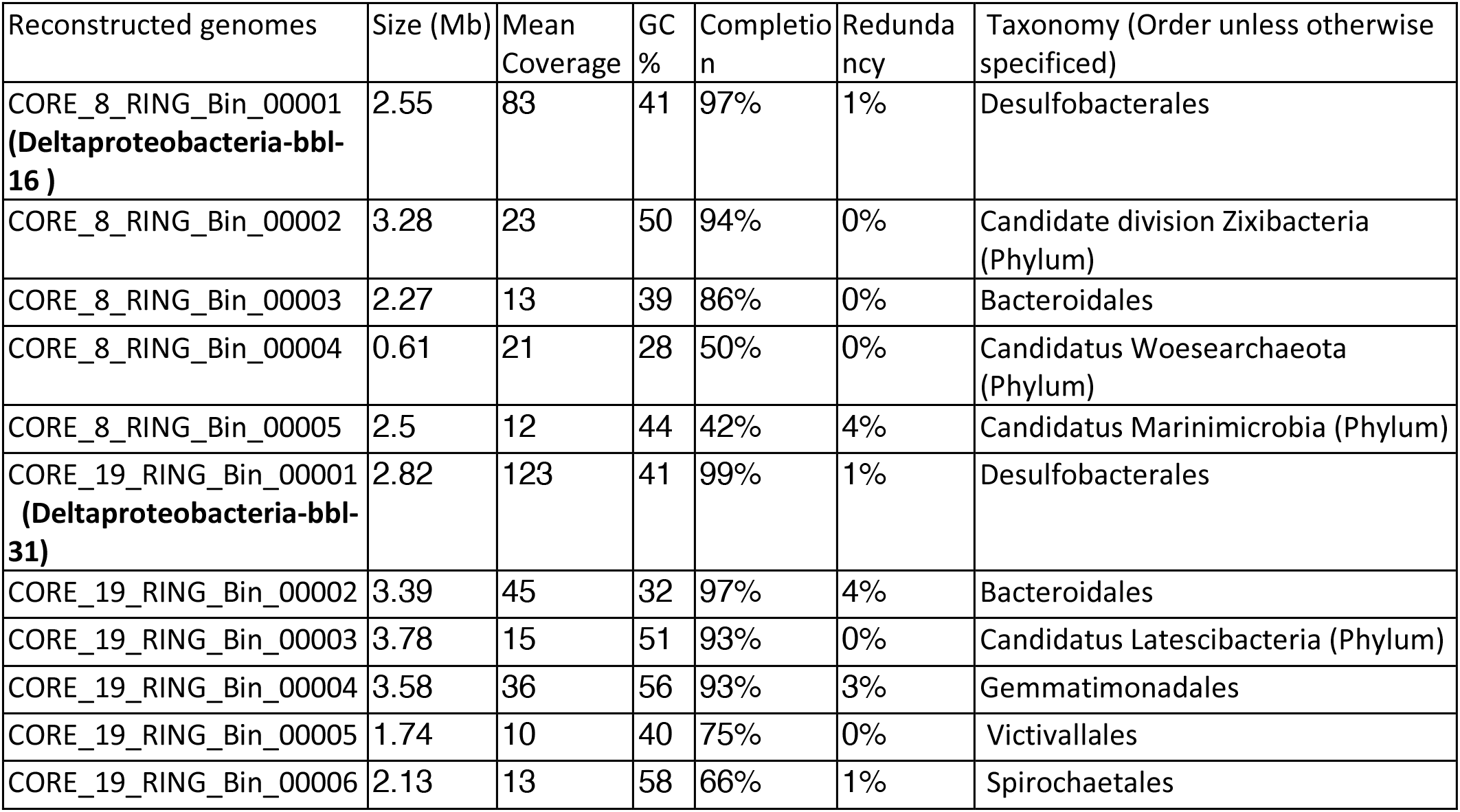
Reconstructed genome statistics including genome size, mean coverage, GC content, percent completion and redundancy, and taxonomic classification.

Consistent with the analysis of diversity, the four sediment samples not associated with a microbial mat resulted in limited assembly as evidenced by metrics such a low N50, low mean contig length, and low total assembly length (**Table S2**), and genome reconstruction was not viable for these samples. In order to assess the relationship between sequencing coverage and diversity in these non-mat sediments, assembly statistics from metagenome sequencing were compared with diversity metrics from the 16S rRNA data (**Table S2**).

### Deltaproteobacteria Dominate Microbial Mats

A near-complete *Deltaproteobacteria* genome most closely related to members of the *Desulfobacula* genus was recovered from the microbial mat at bbl-31. The mean coverage of this genome is 123x, the highest observed from the sample, indicating that this is the most abundant mat organism (**Table S3)**. An 1109 bp fragment of the 16S rRNA recovered from the reconstructed genome aligns with the most abundant 16S rRNA ASV (ASV-1), indicating a consensus between the 16S data set and metagenomic data set, and confirming that the genome of the most abundant organism at the mat was identified and reconstructed. The 16S rRNA sequence has a percent identity of 94.4% with the nearest (*Desulfobacula*) match, indicating that this organism has a marginal affiliation with this genus (a cut-off of ≥94.5% across a full-length sequence would indicate genus-level membership).

The most abundant genome recovered from bbl-16 microbial mat (83x mean coverage) is also a member of *Deltaproteobacteria*, and the average nucleotide identity (ANI) of these two Deltaproteobacteria bbl genomes is 99%. The genomes described here will be referred to as *Deltaproteobacteria-bbl-31* and *Deltaproteobacteria-bbl-16* based on the identity of the proximal waste barrel. In addition to the ANI, the predicted genome size, GC content, and multiple other features of these two genomes are highly similar (**Table S3, S5)**. These findings indicate reproducibility in the microbial response to waste dumping, at two different locations.

### Properties of Dominant Deltaproteobacteria

The near-complete *Deltaproteobacteria-bbl-31* genome size is ∼2.6 Mbps, about half the size of cultivated members of the related *Desulfobacula* genus, which includes several described members: *D. toluolica* (Wöhlbrand et al.), *D. phenolica* (Kuever et al., 2001), and *D*.*TS* (Kim et al., 2014). Additional related Deltaproteobacteria genomes from public databases were identified based on a high percent identity of ribosomal proteins, and are from hydrothermal vent sediments (Dombrowski et al., 2017, 2018) and aquifer groundwater (Castelle et al., 2013). These environmental genomes are similar in size to those from this study and share many of the metabolic traits, discussed in detail in the next section. The genome that has the highest percent identity for the ribosomal proteins is Deltaproteobacteria 457-123 from the Guayamas Basin, and the proteins from this genome and those of Deltaproteobacteria-bbl genomes have a two-way average amino acid identity (AAI) of 52.16% (the AAI genus threshold is 55-60%) (Rodriguez-R and Konstantinidis, 2014) suggesting that these two organisms are related and that both belong to the *Desulfobacteraceae* family.

The metabolic potential for the Deltaproteobacteria-bbl genomes from this study largely overlaps with that of the *Desulfobacula*, and these genomes encode the full pathway for dissimilatory sulfate reduction, including the sulfate adenylyltansferase, adenylylsulfate reductase, and dissimilatory sulfite reductase proteins (**Fig. 2A)**. Similar to members of the *Desulfobacula* genus, these organisms also harbor the capacity for xenobiotic compound degradation, and have the marker gene for anaerobic aromatic degradation, *bamA* (6-oxocyclohex-1-ene carbonyl CoA hydrolase) (Porter and Young, 2013), and the marker gene for reductive dehalogenation (*rdh*) (Hug et al., 2013) (**Fig. 2A)**. Additionally, the full set of required genes for full Benzoyl-CoA to Acetyl-CoA pathway are also encoded, although the metabolic precursor to Benzoyl-CoA remains undetermined in this genome (**Fig. 2A**). These anaerobic aromatic degradation and dehalogenation abilities may be beneficial for this organism in the environment, given the nature of anthropogenic waste (e.g., chlorinated aromatic hydrocarbon compounds) present at this study site (Kivenson et al., 2019).

**Figure 2.**
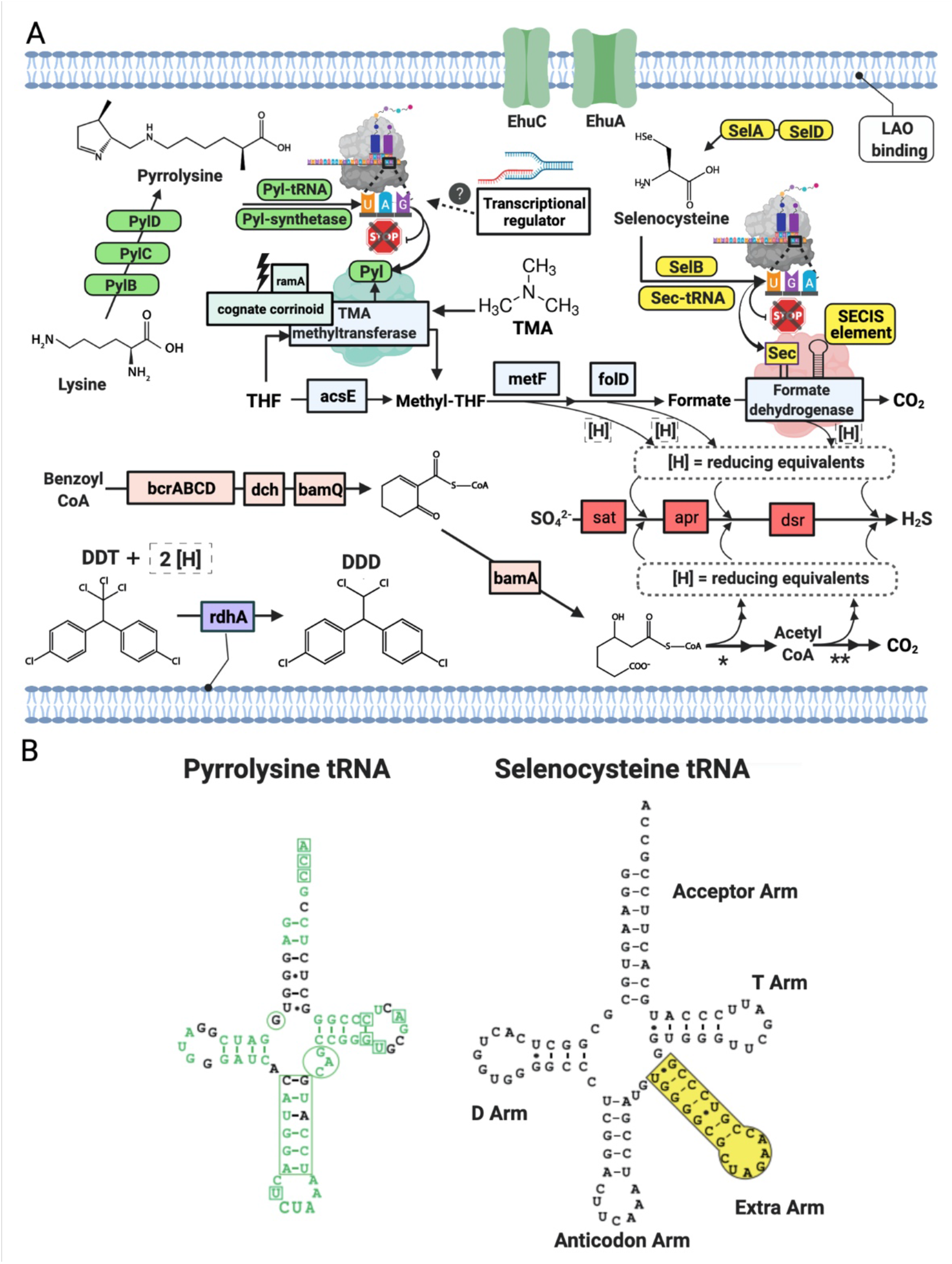
**A)** Schematic of Deltaproteobacterial-bbl cell and metabolic potential with green and yellow icons indicating machinery necessary for genetic code expansion for Pyl and Sec, respectively. Blue icons indicate proteins in the Wood-Ljungdahl pathway, pink icons indicate proteins in the benzoyl CoA pathway, and light purple is involved in dehalogenation. Gene and compound abbreviations: PylB, 3-methylornithine synthase; PylC, 3-methylornithine--L-lysine ligase; PylD, 3-methylornithyl-N6-L-lysine dehydrogenase; Pyl synthetase, pyrrolysyl-tRNA synthetase; Pyl tRNA, pyrrolysine transfer RNA; ramA, methylamine methyltransferase corrinoid protein reductive activase; cognate corrinoid, methyltransferase cognate corrinoid protein; SelD, selenide, water dikinase; SelA, L-seryl-tRNA(Sec) selenium transferase; SelB, selenocysteine-specific translation elongation factor, Sec-tRNA, selenocysteine transfer RNA; SECIS Element, selenocysteine insertion sequence element; EhuC, ectoine hydroxyectoine ABC transporter C; EhuA, ectoine hydroxyectoine ABC transporter A; LAO binding, lysine-arginine-ornithine binding periplasmic protein, rdhA, reductive dehalogenase; DDT, Dichlorodiphenyltrichloroethane; DDD, dichlorodiphenyldichloroethane; THF, tetrahydrofolate; acsE, 5-methyltetrahydrofolate:corrinoid/ iron-sulfur protein co-methyltransferase; metF, methylenetetrahydrofolate reductase; folD, methylene-tetrahydrofolate (CH2-THF) dehydrogenase/cyclohydrolase; bcrABCD, benzoyl CoA reductase subunits A,B,C,D; dch, dienoyl-CoA hydratase; bamQ, 6-hydroxycyclohex-1-ene-1-carbonyl-CoA dehydrogenase; bamA, 6-oxocylcohex-1-ene-1-carbonyl-CoA hydrolase; sat, sulfate adenylyltransferase; apr, adenylylsulfate reductase, dsr, dissimilatory sulfite reductase subunit alpha and beta. **B)** Secondary structures of the pyrrolysine tRNA and selenocysteine tRNA from Deltaproteobacteria-bbl genomes. Green text indicates match with Methanosarcina tRNA sequence, including the corresponding CUA anticodon. Additionally, the boxes and circles show locations of highly conserved regions of the Methanosarcina pyrrolysine tRNA. Small black dots in stems indicate wobble base pairs.

### Metabolic Augmentation by Genetic Code Expansion

Natural genetic code expansion is defined as the ability to encode more than twenty canonical amino acids. The *Deltaproteobacteria bbl-31* genome has the full set of cellular machinery required for both noncanonical amino acids. For Sec utilization in bacteria, a Sec-specific tRNA (*selC*) is required, as well as several proteins: Sec-synthase (alternatively termed L-seryl-tRNA (Sec) selenium transferase; *selA*), a Sec-specific translation elongation factor (*selB*), and selenophosphate synthetase (alternatively termed selenide water dikinase; *selD*, which itself contains a Sec residue) (Stadtman, 1996) (**Fig. 2A**). Along with the Sec-specific machinery proteins, the Sec-tRNA allows the cell to co-translationally insert Sec by repurposing the UGA, or opal codon (which would otherwise signal a stop), in instances when a downstream SECIS element (a ∼50 nt hairpin structure) is present. In the Deltaproteobacteria-bbl genomes (and other Sec-utilizing organisms), the Sec-tRNA differs from most other tRNA: instead of a four-nucleotide variable loop, this tRNA has an extra arm between the anticodon arm and the T arm (**Fig. 2B**).

In addition to the specialized Sec machinery, the incorporation of Sec in the cell is further evidenced by the presence of an in-frame UGA codon followed by the SECIS element in a subset of its genes. In *Deltaproteobacteria-bbl-31*, these features occur in a total of 34 putative selenoproteins, belonging to two groups (**Table S6, S7**). The first group encodes Sec in a conserved position of previously characterized selenoproteins, including those described from marine environments (Zhang et al., 2005; Zhang and Gladyshev, 2007). This group is comprised largely of redox related proteins, and in the Deltaproteobacteria-bbl genome, it includes formate dehydrogenase (**Fig. 2A**), coenzyme F_420_-reducing hydrogenase, methyl-viologen-reducing hydrogenase, sulfurtransferase, a domain of unknown function (DUF 166), Fe-S oxidoreductase, heterodisulphide reductase, a rhodanese-like domain, and the selenophosphate synthetase (**Table S6**). There are several copies of some of these Sec-containing proteins. The second group is comprised of novel, potential selenoproteins, identified using a published *in silico* recognition algorithm (Zhang and Gladyshev, 2005) for locating in-frame UGA residues followed by the SECIS element. These putative new selenoproteins include endonuclease III, cobinamide phosphate guanyltransferase, a histidine kinase, and others (**Table S6**). Although further verification (i.e., confirmation of expression) is needed to confirm the incorporation of Sec and subsequent activity of these proteins, these findings reveal an increased range of proteins in which Sec likely occurs. Furthermore, while Sec is known to increase the catalytic efficiency of proteins, its presence in multiple non-redox proteins suggests that it has additional, as yet unknown role(s).

The rare amino acid, Pyl, is synthesized from two molecules of L-lysine via the action of three proteins: 3-methylornithine synthase (pylB), 3-methylornithine-L-lysine ligase (pylC), and 3-methylornithyl-N6-L-lysine dehydrogenase (pylD) (Gaston et al., 2011b). We analyzed phylogeny and taxonomic diversity for putative PylB protein sequences obtained here, in comparison with other homologs from bacterial and archaeal genomes (**Fig. 3, Fig. S5**). Four monophyletic clades were delineated by taxonomy, but several divergent sequences were separately identified from Firmicutes, Deltaproteobacteria, and Euryarchaeota genomes. Examination of the alignment of all PylB-like sequences revealed conserved regions shared across the four distinct clades (**Fig. 3B**). A radical SAM domain (PF04055) was predicted in most sequences between residues ∼50 to ∼210. All sequences have homology to the pyrrolysine biosynthesis protein PylB family (IPR023891; also homologous to IPR034422 and IPR024021).

**Figure 3.**
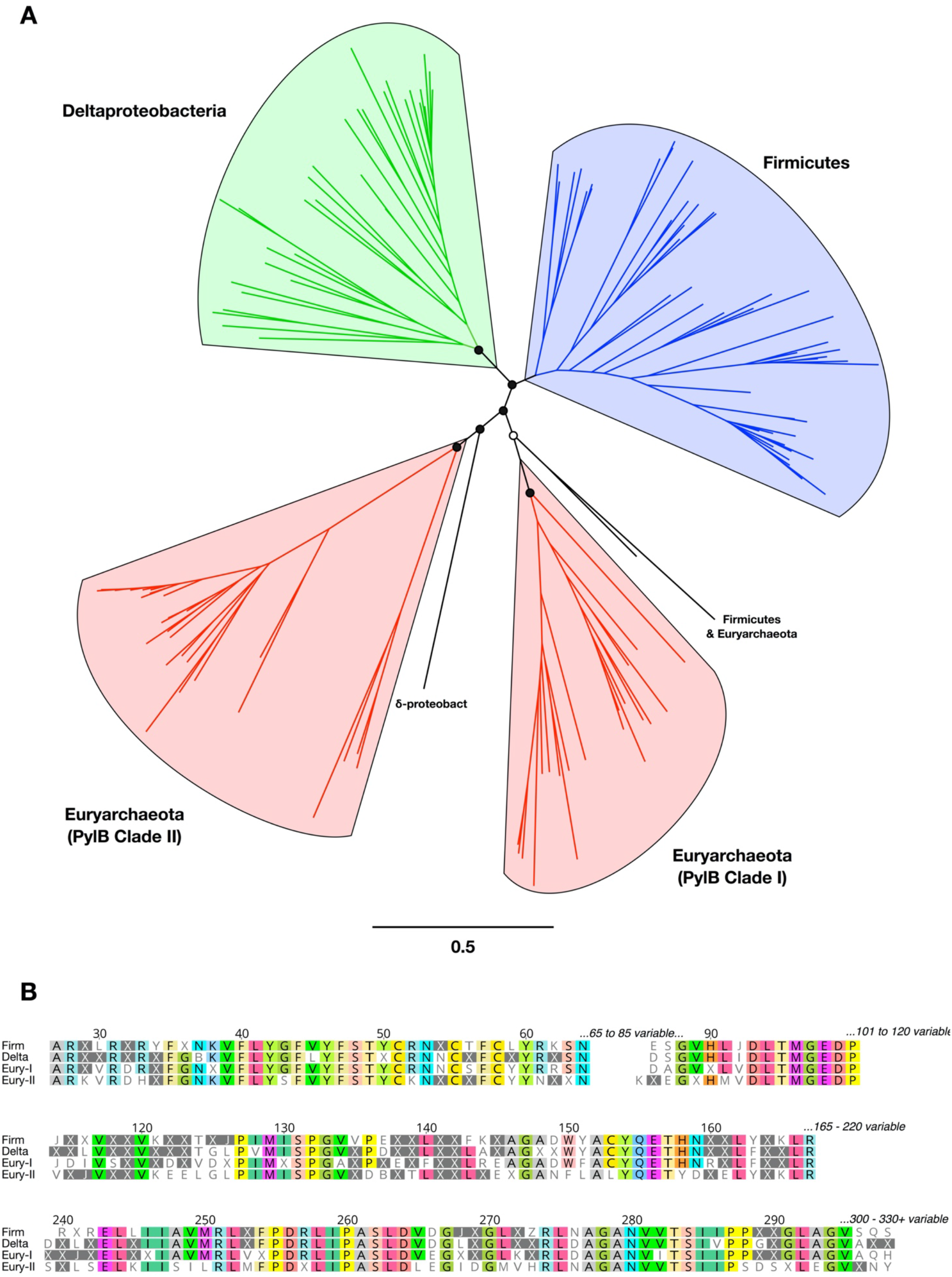
Phylogeny, diversity, and conserved regions of PylB proteins in Bacteria and Archaea. **A)** Unrooted phylogenetic tree from PylB sequence alignment. Major clades are highlighted in color by taxonomic classification. Branch support is shown for the basal clades only (open circles >50%; filled circles >75%). All tip labels and branch support values are provided in Figure S5. **B)** Conserved regions from an alignment of PylB consensus amino acid sequences representing four major clades. A consensus sequence was generated for each of the four PylB clades with a threshold of >50% identical amino acids found in a clade; positions with <50% agreement are indicated with “X”. Conserved positions in the alignment are shown and amino acids are highlighted with a >75% identity threshold.

Co-translational insertion into the growing polypeptide at an in-frame UAG (amber) codon requires Pyl synthetase, and a Pyl-specific tRNA (Srinivasan et al., 2002b; Polycarpo et al., 2004; Gaston et al., 2011b). Along with the Pyl synthesis proteins and synthetase, the specialized amber suppressor tRNA is encoded in the *Deltaproteobacteria-bbl* genomes (**Fig. 2A**). Originally identified in the *Methanosarcina* genus, this Pyl-tRNA has an atypical secondary structure, including a hallmark CUA anticodon, an anticodon stem with an extra nucleotide, and one less nucleotide between the acceptor and D arm in the secondary structure(Srinivasan et al., 2002a). These unusual features are evident in the secondary structure of Pyl-tRNA in the Deltaproteobacteria-bbl genomes, which show a high degree of conservation with the Methanosarcina Pyl-tRNA **(Fig. 2B)**. Earlier studies indicate that incorporation of Pyl is required as part of the active site of mono-, di-, and tri-methylamine methyltransferases(Paul et al., 2000). Both Deltaproteobacteria-bbl genomes have TMA methyltransferase with a conserved active site containing the Pyl residue encoded by an in-frame amber codon at position 327 of 485, (similar to that in archaeal methanogens, which have Pyl at position 334 of 483) (Paul et al., 2000).

In methanogens, the methyl from TMA is transferred by the TMA methyltransferase to a cognate corrinoid protein(Paul et al., 2000). Before accepting the methyl group, the corrinoid protein cobalt (II) active site is reduced to cobalt (I) by the ramA protein, a ferredoxin-like protein with an Fe_4_S_4_ domain (Ferguson et al., 2009); then the methyl group is transferred to coenzyme M reductase (mcrA) and methanogenesis can proceed. With the exception of mcrA, the *Deltaproteobacteria-bbl* genomes have all the aforementioned genes for methyl utilization (**Fig. 2A**); in addition to the Pyl-containing TMA methyltransferase, these organisms have multiple copies of both the cognate corrinoid protein and ramA. Additionally, the Deltaproteobacteria-bbl genomes contain three copies of the gene for methionine synthase (metH), which is homologous to the methylotrophic methanogenesis methyltransferase (Paul and Krzycki, 1996), and the chloromethane utilization methyltransferase (mtaA-cmuA), which belongs to a protein family with high similarity to methyltransferases of methanogenic archaea(Vannelli et al., 1999).

Rather than transferring the methyl group from TMA to mcrA as occurs for methanogenesis, an alternate path may be utilized, wherein the methyl group is transferred to tetrahydrofolate and enters the Wood Ljungdahl (WL) pathway, for which the full set of proteins is encoded in the *Deltaproteobacteria-bbl* genomes (**Fig. 2A)**. The WL pathway operates in reverse (for carbon compound oxidation) in *Desulfobacula* and other sulfate reducing bacteria (Wöhlbrand et al.), and may be paired with sulfate reduction, and ultimately used for energy and / or carbon assimilation for the cell. Intriguingly, two of the key proteins in the proposed reverse WL pathway contain residues arising from independent instances of genetic code expansion: the Pyl-containing TMA methyltransferase and the Sec-containing formate dehydrogenase (**Fig. 2A)**. This dual genetic code expansion-augmented metabolism may be beneficial for the cell due to linking of advantageous traits: the utilization of a ubiquitously-available compound (TMA) by the Pyl-containing methyltransferase, and the increased catalytic efficiency conferred by the Sec-containing formate dehydrogenase.

Additional evidence for the importance of genetic code expansion-augmented metabolism in the Deltaproteobacteria-bbl (and other Deltaproteobacterial) genomes is the presence of multiple genes relevant to Pyl synthesis and methyl amine metabolism, proximal to the TMA methyltransferase gene (**Table S8**). This functional gene organization may have biological significance, as it is completely conserved within the Deltaproteobacteria-bbl genomes, and is highly similar in related genomes from other locations (discussed further in the following section on biogeographic distribution). In the *Deltaproteobacteria-bbl* and related genomes, the gene adjacent to TMA methyltransferase encodes a lysine-arginine-ornithine binding periplasmic protein, which may play a role in Pyl synthesis, as lysine is the precursor to pyrrolysine (Krzycki, 2013). Moreover, the addition of ornithine to growth media has been shown to increase the yield of full-length Pyl proteins (Namy et al., 2007). The following two genes encode ectoine / hydroxyectoine permeases / ABC transporters; ectoine and hydroxyectoine are compatible solutes and the aforementioned transporter class of proteins is involved in the transport of methylated amines into the cell (Burke et al., 1998). Also proximal to the methyltransferase gene is a transcriptional regulator, though a more specific functional role remains undetermined. In members of the *Firmicutes*, transcriptional feedback loops control the genetic code expansion and subsequent use of TMA via the TMA methyltransferase(Prat et al., 2012). A glycine betaine methyltransferase gene also occurs in this locus, which is commonly mis-annotated as a trimethylamine methyltransferase (Ticak et al., 2014), but does not contain pyrrolysine. In other organisms, the role of this protein is to convert glycine betaine to dimethylglycine and methylcobalamin (Ticak et al., 2014). Several additional copies of the nonpyrrolysine methyltransferase are present elsewhere in each of the *Deltaproteobacteria-bbl* and related genomes, suggestive of a broad capacity for methyl metabolism in these organisms.

### Deltaproteobacteria Compete with Methanogens for TMA

Based on the capacity for TMA methyl-group oxidation encoded by the *Deltaproteobacteria-bbl-31* and *Deltaprotebacteria-bbl-16*, we mined published metagenomic and metatranscriptomic data from sulfidic sediments for indications of Pyl incorporation by *Deltaproteobacteria* (**Table S9**). In reanalyzing a recent study of Baltic Sea sediment (Thureborn et al., 2016) we identified *Deltaproteobacterial* expression of the TMA methyltransferase, and of the Pyl synthetase, PylD and PylC proteins (**Fig. 4**). Furthermore, the expressed TMA methyltransferase from the Baltic Sea transcriptomic dataset is nearly identical (96%) with that of the Deltaproteobacterial-bbl protein. Importantly, the position of the pyrrolysine is conserved, as well as the amino acid sequence following the UAG codon (**Fig. 5A**).

**Figure 4.**
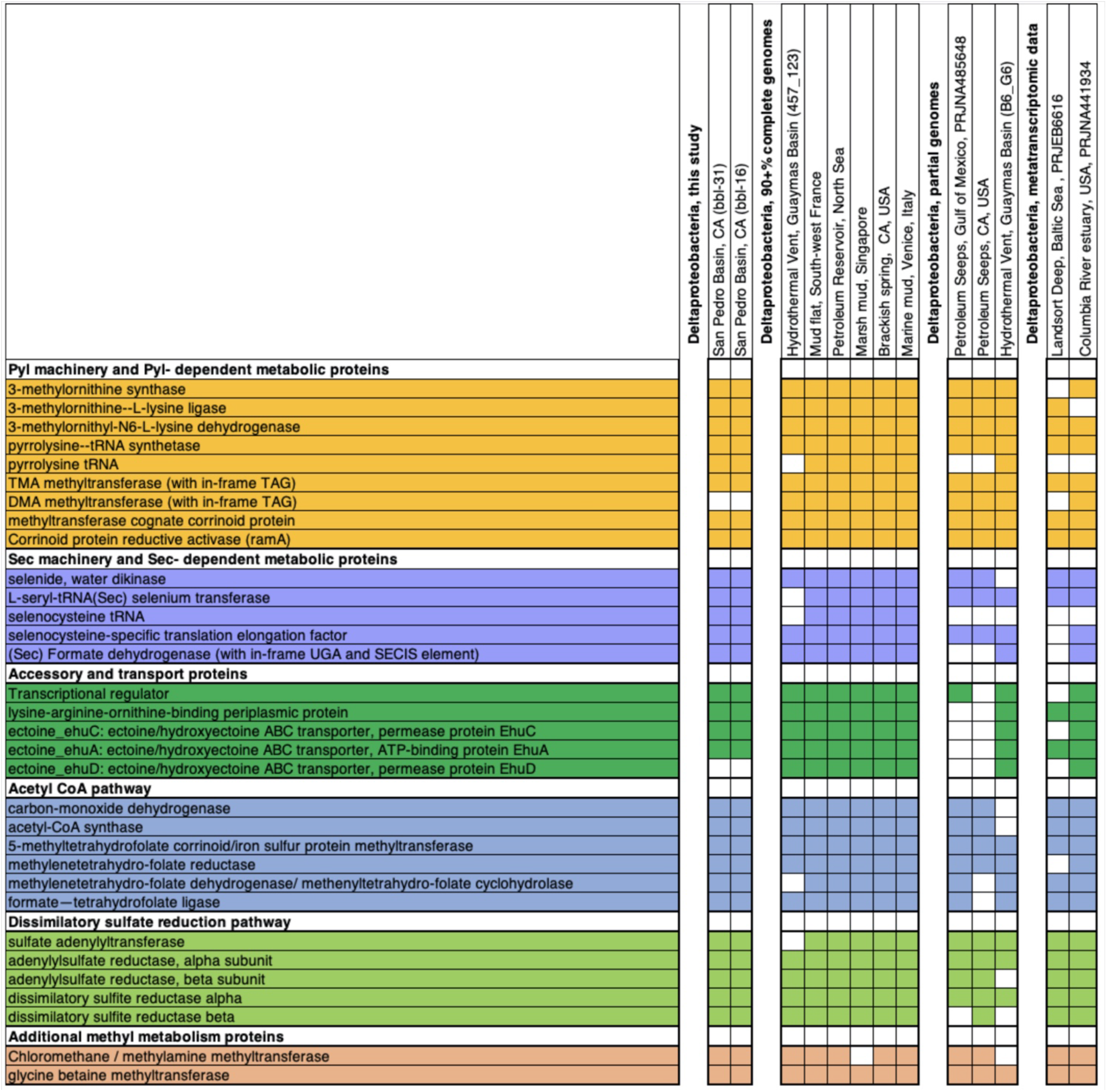
Genes from metabolic, biosynthetic, and other pathways related to genetic code expansion-enabled methyl utilization, shown from Deltaproteobacteria from this study along with other genomic and transcriptomic data sets. Colored boxes indicate that the gene is present in Deltaproteobacteria as either transcriptomic or genomic sequence.

**Figure 5.**
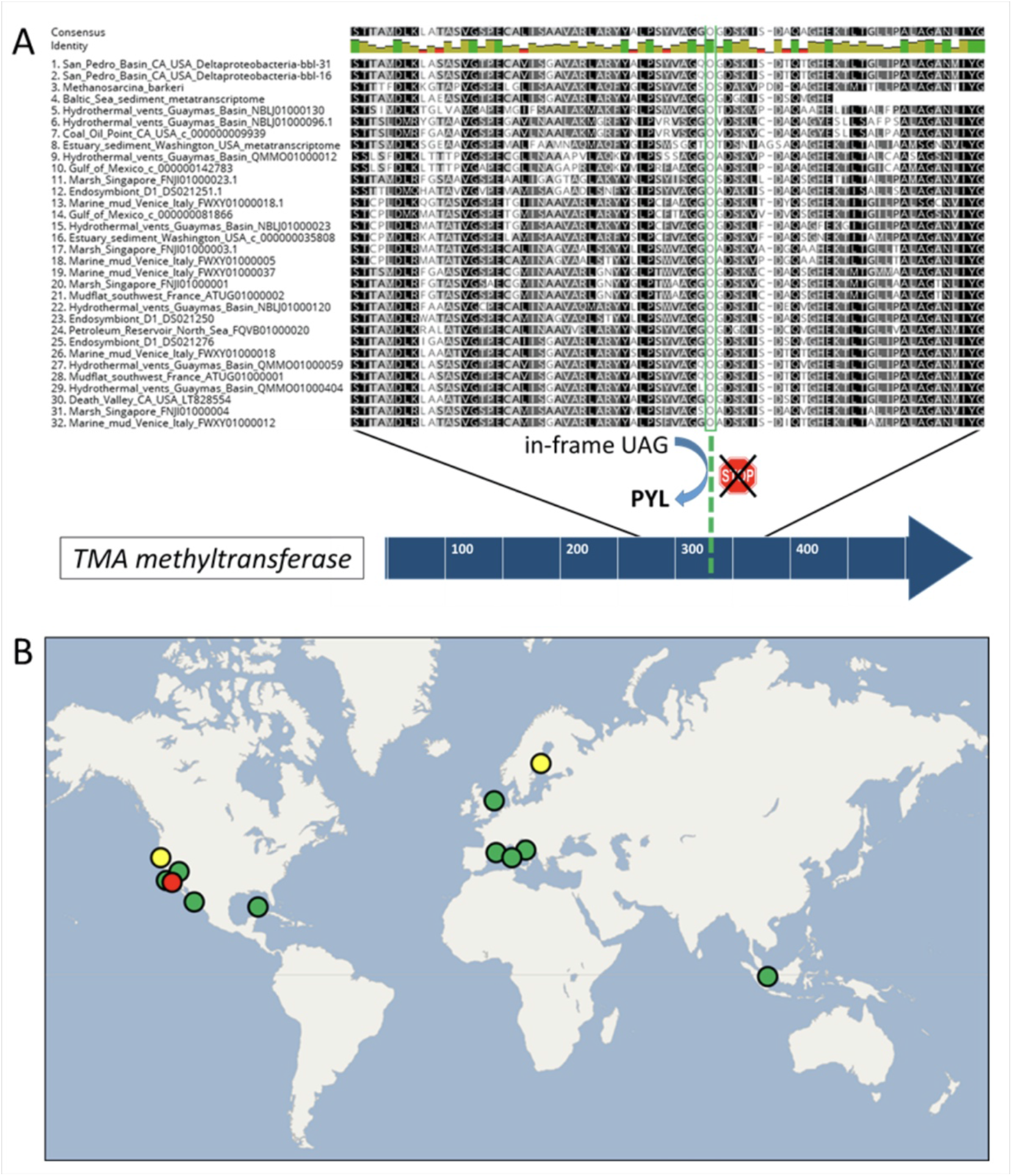
**A)** Alignment of select residues of the trimethylamine methyltransferase protein, spanning the pyrrolysine residue (in green), as well as the conserved residues following the UAG codon. Proteins are shown from the Deltaproteobacteria-bbl genomes from this study, Methanosarcina barkeri for comparison, and Deltaproteobacteria from other studies, including two transcriptomics data sets. **B)** The global distribution of dual genetic code expansion-enabled Deltaproteobacteria that have the trimethylamine methyltransferase and conserved Pyl residue, as well as Sec and Pyl machinery genes. Circle colors are as follows: red indicates this study, yellow indicates metatranscriptomic data sets, and green includes metagenomic / genomic data sets. The location source and type of each dataset is provided in Table S9.

These observations reveal that TMA metabolism enabled by genetic code expansion is actively utilized by *Deltaproteobacteria* in the environment, putting them in competition with methylotrophic methanogens. While methanogens (primarily *Methanosarcina*) are active at the Baltic Sea site, there is no detectable expression of TMA (or DMA or MMA) –methyltransferase by any archaea. The primary expressed methanogenesis pathway is hydrogenotrophic (CO_2_+ H_2_), indicating that oxidation of TMA by *Deltaproteobacteria* may be an important environmental fate for this compound. Analysis of an additional sediment metatranscriptome from the Columbia River estuary, Washington, USA shows the expression of Deltaproteobacterial Pyl ligase, PylB, PylC, and TMA methyltransferase spanning the Pyl-containing active site, and Deltaproteobacterial expression of Sec machinery proteins and Sec-containing formate dehydrogenase and other one-carbon metabolic pathway proteins (**Fig. 4**).

While TMA has been referred to as a non-competitive substrate that is used in anoxic sediments exclusively by methanogenic archaea, for example (Reeburgh, 2007; Joye et al., 2009; Orsi et al., 2013; Kelley et al., 2014; Zhuang et al., 2016), the pioneering studies that first examined methylated amine utilization in anoxic sediments demonstrated that TMA consumption rates exceed their methanogenic potential and that bacteria also utilize methyl amines (King, 1984a, 1984b). Moreover, addition of sulfate reduces the fraction of substrate metabolized to methane from trimethylamine in incubation experiments(Lovley and Klug, 1983). Similarly, a more recent study of TMA utilization in marine sediments demonstrated that inhibition of sulfate reducing bacteria increased the rate of TMA-methanogenesis(Xiao et al., 2018), and another study describes that TMA-amended incubations resulted in H_2_S production (Cadena et al., 2018), again pointing to competitive TMA utilization by anaerobic bacteria. Collectively, available genomic, transcriptomic and environmental data indicate that Deltaproteobacteria compete with methanogens for TMA, using metabolism dependent on genetic code expansion.

### A Cosmopolitan Distribution for Deltaproteobacteria with Genetic Code Expansion-Augmented Metabolism

To assess the geographic distribution of *Deltaproteobacteria* with metabolism augmented by genetic code expansion, we surveyed publicly-available Deltaproteobacteria genomes from marine and terrestrial environments for GCE genes (**Table S9, S10**). In addition to the genomes from this study, nine other *Deltaproteobacterial* genomes (six of which are ≥90% complete) were identified with the Pyl and Sec biosynthesis proteins (**Fig. 4)**. The Deltaproteobacteria from these studies also encode a TMA methyltransferase protein with a conserved Pyl residue (**Fig. 5A**). These bacteria originate from diverse settings; in addition to the San Pedro Basin (this study), they include hydrothermal vents in the Guaymas Basin, marine mud in Italy, petroleum seeps in the North Sea and California, as well as terrestrial environments, including a marsh in Singapore and brackish spring in California (**Fig. 5B**). Most of these Deltaproteobacteria have multiple copies of TMA methyltransferase (with up to five copies in a single genome) and the dimethylamine methyltransferase with an in-frame Pyl. Additionally, most of the genomes have all of the proteins necessary for the reverse WL pathway, including a Sec-containing formate dehydrogenase (**Fig. 4**). These specialized bacteria inhabit many settings that are oxygen-limited and hydrocarbon-rich, much like the related *Desulfobacula* genus. Taken together, the distribution of these Deltaproteobacteria and their genomic repertoire indicates an important geochemical role governing the fate for TMA in the environment.

## Methods

### Site Description and Sampling

The study site is located at ∼894 meter water depth in the oxygen-limited San Pedro Basin, off the coast of California, at 33.566N, 118.426W. It was accessed in October 2013 using the autonomous underwater vehicle (AUV) *Sentry* (dives S-202 and S-208) and the remotely operated vehicle (ROV) *Jason* (dive J2-746), during RV *Atlantis* Leg 26-06, as previously described (Valentine et al., 2016; Kivenson et al., 2019). Briefly, the *AUV* Sentry was deployed to locate the industrial waste dumpsite, and high-resolution multibeam echosounder bathymetry at 60 m above the seafloor indicated the presence of positive elevation anomalies consistent with barrels; a photo-survey was conducted, showing multiple barrels on the seafloor. The *ROV* Jason was then deployed for additional images of the barrels, and to collect sediment samples for chemical measurements (previously described (Kivenson et al., 2019)), and biological analyses. The core samples used for biological analyses include the surficial (top two centimeters) sediment from two push cores not affiliated with barrels (collected >60m from a barrel), two microbial mat/ ring samples (collected at the microbial ring-shaped mat from each of two barrels), and two samples from just outside the microbial ring/mat (<0.25m). Sediment samples for biological analyses were sectioned shipboard following *ROV* retrieval, and stored at −80°C prior to DNA extraction.

### DNA extraction and preparation

Six core-top (0-2cm) sediment samples were processed for 16S rRNA and metagenomic sequencing. First, bulk DNA was extracted from ∼0.25 grams of each sediment sample using a MoBio Powersoil DNA extraction kit according to manufacturer instructions, with the following modifications: bead beating (60 sec.) to enhance cellular lysis for greater DNA yield, and a secondary purification step using a Qiagen AMPure XP bead cleanup kit to remove humics and other organic matter.

### 16S rRNA sequencing and analyses

The V4 region of the 16S SSU rRNA gene was amplified using primers (515F-806R) optimized for sequencing on the Illumina MiSeq platform as described previously (Kozich et al., 2013; Apprill et al., 2015). Amplicon PCR reactions contained 1µl template DNA (5ng/µl), 2µl forward primer, 2µl reverse primer, and 17µl AccuPrime Pfx SuperMix. Thermocycling consisted of 95°C for 2min, 30 cycles of 95°C for 20s, 55°C for 15s, 72°C for 5min, and a final elongation at 72°C for 10min. Sample concentrations were normalized using the SequelPrep Normalization Kit and visualized on an Agilent 2200 Tapestation with the genomic DNA ScreenTape at the California NanoSystems Institute at UC Santa Barbara. Samples were sequenced at the UC Davis Genome Center on the Illumina MiSeq platform with 250nt, paired end reads. Raw sequences were quality checked and analyzed using the two open-source software packages: mothur v.1.36.1 for operational taxonomic units (OTUs) (Kozich et al., 2013) and DADA2 v.1.1.0 for amplicon sequence variants (ASVs) (Callahan et al., 2016, 2), using the SILVA bacterial reference database (Release 128) (Pruesse et al., 2007). The standard Miseq SOP pipeline using mothur (Kozich et al., 2013) was performed to enable comparison with another San Pedro Basin dataset referenced in the results, and the ASV pipeline using DADA2 was used in order to resolve fine scale sequence variation (Callahan et al., 2016, 2). Phyloseq was used to for visualization, to examine community diversity and ASV abundance (McMurdie and Holmes, 2013).

### Metagenomic preparation and sequencing

For metagenomic sequencing, following purification, DNA concentrations were determined using Invitrogen Qubit fluorometric quantification and normalized for library preparation (5ng/µL). The Nextera XT library preparation kit was used in accordance with manufacturer instructions with custom Illumina indexes for demultiplexing. The concentration and size of each sample following library preparation were validated using the Qubit and Agilent 2200 Tapestation with the genomic DNA ScreenTape, respectively. Whole genome sequencing was performed at the California NanoSystems Institute at UC Santa Barbara using the Illumina NextSeq 500 platform, using v2 chemistry and the Mid Output flow cell kit, generating 78.8 GB of paired end 150 bp reads representing the collected sampled DNA.

### Genome reconstruction from whole genome sequencing data

Quality control for reads was performed using Trimmomatic v.0.36 to remove sequencing artifacts and low quality reads (Bolger et al., 2014). Adapters and short reads were removed using the adaptive read trimmer, Sickle v.1.33 (Joshi and Fass, 2018). After quality control, reads for each sample were individually assembled using MetaSpades v.3.10.1. Additionally, Megahit v.1.1.2 assembly was generated for comparison and included in Table S2 (Li et al., 2015). Default parameters were used for all software unless otherwise specified. The quality of the assemblies was determined using QUAST v.4.6 (Gurevich et al., 2013). Trimmed (post quality-control) sequencing reads for each sample were mapped to assembled contigs ≥ 1000 bp, using Bowtie2 v.2.3 and Samtools v.1.7 (Langmead et al., 2009; Li et al., 2009). GC content, tetranucleotide content, and coverage were used manually within Anvi’o v.6.2.0 to generate genome bins (Eren et al., 2015). The bins were manually refined using Anvi’o on the basis of consistency of coverage and nucleotide content of contigs in each bin, in order to reconstruct high-quality metagenome-assembled genomes (MAGs). Completion and redundancy for each refined MAG was determined with Anvi’o on the basis of recovery (completion) and repeats (redundancy) of conserved single copy genes of the nearest taxonomic group.

### Functional annotation and taxonomic assignment of genomes

Predicted proteins in each genome were identified using Prodigal v.2.6.3 individually (single genome mode) and functional annotation was performed using Hmmer v.3.1, with hmmscan (1E-10) with top hits only, against the Pfam database v.31 and Tigrfam database v.15.0 (Haft et al., 2003; Hyatt et al., 2010; Finn et al., 2011, 2013). Additional protein annotation was performed to assign proteins to COG categories using the rps-blast option in Blast v.2.7.1 (Altschul et al., 1990) and the Perl script cdd2cog, available at https://github.com/aleimba/bac-genomics-scripts/tree/master. RNAmmer v.1.2 was used for identification and recovery of 16S rRNA sequences from the reconstructed genomes (Lagesen et al., 2007). Taxonomy of reconstructed genomes was determined with preference given to criteria in the following order: (i) 16S rRNA genes, when recovered, (ii) recA, due to its strong phylogenetic signal, and (iii) taxonomic assignment using other ribosomal protein sequences (Rp_S8) (Table 1, Table S3). GTDB-TK v.1.3.0 was used to assign taxonomic identification using the RS95 database release, and is included for reference (Table S4) (Chaumeil et al., 2020), but established taxonomic nomenclature is used throughout the manuscript for consistency. Recovered genome-origin 16S sequences were aligned using Muscle v.3.8 with candidate ASV sequences from the 16S rRNA data set to determine the percent abundance of corresponding genomes (Edgar, 2004). To compare genomes from different samples, the average nucleotide identity (ANI) and average amino acid identity (AAI) were used to measure nucleotide level genomic similarity for genomes with high completion levels, using the Enveomics webserver ANI and AAI calculators (Rodriguez-R and Konstantinidis, 2016).

### Characterization of GCE and application of noncanonical protein translation tables

To identify instances of genetic code expansion in reconstructed genomes, the functional annotation results for genomes were compared with previously described cellular machinery required for the co-translational insertion of Sec and Pyl (Table S10). Additionally, the requisite Sec- and Pyl-specific tRNA sequences, as well as their secondary structures and corresponding anticodons were identified using the Aragorn web server (Laslett and Canback, 2004). The anticodon CTA was identified for TAG stop-codon suppression and Pyl insertion, and the anticodon TCA was identified for TGA stop-codon suppression and Sec insertion, as well as the elongated extra arm of the Sec tRNA.The Sec-tRNA structure was further validated using the Secmarker tool via the webserver at https://secmarker.crg.es/ and Pyl-tRNA structure was confirmed using Infernal v.1.1.2 (Nawrocki and Eddy, 2013; Santesmasses et al., 2017). The secondary structures of each tRNA were also manually inspected against previously characterized Sec- and Pyl-tRNA structures.

For genomes with an expanded genetic code, alternate protein prediction methods were applied for identification and validation of repurposed TAG and TGA codons. For Pyl residue identification in proteins, the source code for Prodigal was modified to implement a custom translation table with in-frame TAG readthrough, as previously described (Kivenson and Giovannoni, 2020). The modified code and documentation for the TAG readthrough translation table is freely available at https://github.com/VeronikaKivenson/Prodigal. Next, protein sequence alignment was performed for proteins with in-frame TAG codons in the relevant methyltransferases using Muscle v.3.8 (Edgar, 2004). These alignments were then manually inspected to determine if the region containing and following the TAG codon in Deltaproteobacterial proteins is conserved as compared to pyrrolysine-containing methyltransferase protein sequences originating from archaea (Paul et al., 2000).

Proteins containing selenocysteine were identified and validated using a list of previously characterized selenoproteins (Zhang and Gladyshev, 2005, 2007, 2008; Zhang et al., 2005; Peng et al., 2016). These proteins as well as additional, novel selenoproteins were identified and analyzed using the program, bSECISearch, which applies a bacterial SECIS consensus model for the detection and identification of the SECIS element and the in-frame UGA sequence encoding Sec (Zhang and Gladyshev, 2005).

For PylB phylogenetic reconstruction, the MAG-derived proteins from this study were used to search for homologs, from which a non-redundant set of representatives was generated using CD-HIT (Li and Godzik, 2006), with a 90% global identity threshold. Next, sequences were aligned using MUSCLE (Edgar, 2004) and PylB protein trees were constructed using FastTree v.2.1.5 (Price et al., 2010) with the Whelan and Goldman model and optimized Gamma20 likelihood. For comparison, another PylB tree was constructed using RAXML v.8.2.12 (Kozlov et al., 2019), with the LG substitution matrix, and automatic determination of optimal bootstraps (**Fig. S6, S7**).

### Identification and distribution of specialized Deltaproteobacteria in the environment

Publicly available data sets were examined to determine the environmental distribution and niche preference of specialized Deltaproteobacteria with an expanded genetic code and augmented metabolism. A list of candidate GCE-capable Deltaproteobacterial genomes was compiled on the premise of ribosomal protein sequence similarity with *Deltaproteobacterial-bbl* genomes identified in this study (Table S9). Annotation of these candidate genomes was performed to determine if they encode the requisite components for GCE, as described in the preceding section.

Second, short read datasets from select studies on oxygen-limited environments were analyzed due to their ability to host the most closely related and well-characterized genera: the *Desulfobacula*. The targeted locations included hydrothermal vents, oil seeps, anaerobic sediments, and other oxygen-limited environments. Sequencing reads were accessed from the NCBI SRA site on June 19, 2019, and downloaded using the SRA-dump option of the NCBI SRA Toolkit v2.8.1. These short-read data sets were individually assembled, and annotation of the resulting contigs and partial genomes was performed as previously described. Taxonomic assignment was determined using ribosomal and other protein sequences to identify and confirm Deltaproteobacterial contigs and partial genomes, and subsequent identification of GCE machinery and augmented metabolism was performed as previously described.

Third, protein searches were performed using the amino acid sequences of Deltaprotebacteria-bbl proteins that are central to GCE and augmented metabolism pathways from this study. The JGI IMG webserver (https://gold.jgi.doe.gov) was accessed on July 30, 2019 for data from the Genomes Online Database (GOLD) resource (Mukherjee et al., 2021). This data further supplemented the data examined in the previous two steps, and led to the identification and analysis of metatranscriptomic data sets from sediments from geographic areas of interest. Following quality control, the assembly, protein prediction, and gene annotation steps were performed as previously described, with the addition of a secondary protein prediction step using FragGeneScan v.1.30 for identification of proteins in fragmented sequences in metatranscriptomic data (Rho et al., 2010).

Chemdraw and Biorender software were used for creating graphics for primary and supplementary figures. The computational biology data processing and analysis workflow was completed using the Extreme Science and Engineering Discovery Environment (XSEDE) Bridges resource at the Pittsburgh Supercomputing Center (Towns et al., 2014; Nystrom et al., 2015).

## Author Contributions

VK, BGP, and DLV designed the investigation and collected the samples. VK prepared samples for sequencing, performed 16S rRNA and metagenome processing, metabolic reconstructions and identified re-purposed stop codons. VK and BGP identified and analyzed genetic code expansion machinery. All authors contributed to manuscript preparation.

## Acknowledgments

We thank the captain and crew of the RV Atlantis, the pilots and crew of the ROV Jason, the crew of the AUV Sentry, and the scientific party of the AT-26-06 expedition. We thank Stephen J. Giovannoni, A. Murat Eren, and Tom O. Delmont for helpful discussions and metagenomic analysis support.

## Data Availability Statement

Sequence data will be deposited in the NCBI SRA under BioProject # following manuscript submission. Metagenome-assembled genomes will be made available under NCBI BioProject #. Modification of Prodigal software and supporting documentation is available at https://github.com/VeronikaKivenson/Prodigal.

## References

Altschul, S. F., Gish, W., Miller, W., Myers, E. W., and Lipman, D. J. (1990). Basic local alignment search tool. J. Mol. Biol. 215, 403–410. doi:10.1016/S0022-2836(05)80360-2.

Apprill, A., McNally, S., Parsons, R., and Weber, L. (2015). Minor revision to V4 region SSU rRNA 806R gene primer greatly increases detection of SAR11 bacterioplankton. Aquatic Microbial Ecology 75, 129–137.

Atkins, J. F., and Gesteland, R. (2002). The 22nd Amino Acid. Science 296, 1409–1410. doi:10.1126/science.1073339.

Axley, M. J., Böck, A., and Stadtman, T. C. (1991). Catalytic properties of an Escherichia coli formate dehydrogenase mutant in which sulfur replaces selenium. PNAS 88, 8450–8454. doi:10.1073/pnas.88.19.8450.

Böck, A. (2000). Biosynthesis of selenoproteins—an overview. Biofactors 11, 77–78.

Böck, A., Forchhammer, K., Heider, J., Leinfelder, W., Sawers, G., Veprek, B., et al. (1991). Selenocysteine: the 21st amino acid. Molecular Microbiology 5, 515–520. doi:10.1111/j.1365-2958.1991.tb00722.x.

Bolger, A. M., Lohse, M., and Usadel, B. (2014). Trimmomatic: a flexible trimmer for Illumina sequence data. Bioinformatics 30, 2114–2120. doi:10.1093/bioinformatics/btu170.

Borges, A. V., Champenois, W., Gypens, N., Delille, B., and Harlay, J. (2016). Massive marine methane emissions from near-shore shallow coastal areas. Scientific Reports 6, 27908. doi:10.1038/srep27908.

Burke, S. A., Lo, S. L., and Krzycki, J. A. (1998). Clustered Genes Encoding the Methyltransferases of Methanogenesis from Monomethylamine. Journal of Bacteriology 180, 3432–3440.

Cadena, S., García-Maldonado, J. Q., López-Lozano, N. E., and Cervantes, F. J. (2018). Methanogenic and Sulfate-Reducing Activities in a Hypersaline Microbial Mat and Associated Microbial Diversity. Microbial Ecology 75, 930–940. doi:10.1007/s00248-017-1104-x.

Callahan, B. J., McMurdie, P. J., Rosen, M. J., Han, A. W., Johnson, A. J. A., and Holmes, S. P. (2016). DADA2: High-resolution sample inference from Illumina amplicon data. Nature Methods 13, 581–583. doi:10.1038/nmeth.3869.

Castelle, C. J., Hug, L. A., Wrighton, K. C., Thomas, B. C., Williams, K. H., Wu, D., et al. (2013). Extraordinary phylogenetic diversity and metabolic versatility in aquifer sediment. Nature communications 4, 2120.

Chaumeil, P.-A., Mussig, A. J., Hugenholtz, P., and Parks, D. H. (2020). GTDB-Tk: a toolkit to classify genomes with the Genome Taxonomy Database. Bioinformatics 36, 1925–1927. doi:10.1093/bioinformatics/btz848.

Chen, Y. (2012). Comparative genomics of methylated amine utilization by marine Roseobacter clade bacteria and development of functional gene markers (tmm, gmaS). Environmental Microbiology 14, 2308–2322. doi:10.1111/j.1462-2920.2012.02765.x.

Cone, J. E., Del Río, R. M., Davis, J. N., and Stadtman, T. C. (1976). Chemical characterization of the selenoprotein component of clostridial glycine reductase: identification of selenocysteine as the organoselenium moiety. Proc. Natl. Acad. Sci. U.S.A. 73, 2659–2663.

Copeland, P. R. (2005). Making sense of nonsense: the evolution of selenocysteine usage in proteins. Genome Biol 6, 221. doi:10.1186/gb-2005-6-6-221.

Council on Environmental Quality (1970). Ocean Dumping: A National Policy. A Report to the President. US Government Printing Office.

Dombrowski, N., Seitz, K. W., Teske, A. P., and Baker, B. J. (2017). Genomic insights into potential interdependencies in microbial hydrocarbon and nutrient cycling in hydrothermal sediments. Microbiome 5, 106. doi:10.1186/s40168-017-0322-2.

Dombrowski, N., Teske, A. P., and Baker, B. J. (2018). Expansive microbial metabolic versatility and biodiversity in dynamic Guaymas Basin hydrothermal sediments. Nature Communications 9, 4999. doi:10.1038/s41467-018-07418-0.

Duedall, I. W., Ketchum, B. H., Park, P. K., and Kester, D. R. (1983). Global inputs, characteristics, and fates of ocean-dumped industrial and sewage wastes: an overview. New York, New York: John Wiley & Sons.

Edgar, R. C. (2004). MUSCLE: multiple sequence alignment with high accuracy and high throughput. Nucleic Acids Res 32, 1792–1797. doi:10.1093/nar/gkh340.

Eren, A. M., Esen, Ö. C., Quince, C., Vineis, J. H., Morrison, H. G., Sogin, M. L., et al. (2015). Anvi’o: an advanced analysis and visualization platform for ‘omics data. PeerJ 3, e1319.

Ferguson, T., Soares, J. A., Lienard, T., Gottschalk, G., and Krzycki, J. A. (2009). RamA, a Protein Required for Reductive Activation of Corrinoid-dependent Methylamine Methyltransferase Reactions in Methanogenic Archaea. J. Biol. Chem. 284, 2285–2295. doi:10.1074/jbc.M807392200.

Finn, R. D., Bateman, A., Clements, J., Coggill, P., Eberhardt, R. Y., Eddy, S. R., et al. (2013). Pfam: the protein families database. Nucleic acids research 42, D222–D230.

Finn, R. D., Clements, J., and Eddy, S. R. (2011). HMMER web server: interactive sequence similarity searching. Nucleic acids research 39, W29–W37.

Gaston, M. A., Jiang, R., and Krzycki, J. A. (2011a). Functional context, biosynthesis, and genetic encoding of pyrrolysine. Current opinion in microbiology 14, 342–349.

Gaston, M. A., Zhang, L., Green-Church, K. B., and Krzycki, J. A. (2011b). The complete biosynthesis of the genetically encoded amino acid pyrrolysine from lysine. Nature 471, 647– 650. doi:10.1038/nature09918.

Gurevich, A., Saveliev, V., Vyahhi, N., and Tesler, G. (2013). QUAST: quality assessment tool for genome assemblies. Bioinformatics 29, 1072–1075.

Haft, D. H., Selengut, J. D., and White, O. (2003). The TIGRFAMs database of protein families. Nucleic acids research 31, 371–373.

Hug, L. A., Maphosa, F., Leys, D., Löffler, F. E., Smidt, H., Edwards, E. A., et al. (2013). Overview of organohalide-respiring bacteria and a proposal for a classification system for reductive dehalogenases. Phil. Trans. R. Soc. B 368, 20120322.

Hyatt, D., Chen, G.-L., LoCascio, P. F., Land, M. L., Larimer, F. W., and Hauser, L. J. (2010). Prodigal: prokaryotic gene recognition and translation initiation site identification. BMC bioinformatics 11, 119.

Joshi, N., and Fass, J. (2018). (2011). Sickle: A sliding-window, adaptive, quality-based trimming tool for FastQ files. Available at: https://github.com/najoshi/sickle [Accessed July 10, 2018].

Joye, S. B., Samarkin, V. A., Orcutt, B. N., MacDonald, I. R., Hinrichs, K.-U., Elvert, M., et al. (2009). Metabolic variability in seafloor brines revealed by carbon and sulphur dynamics. Nature Geoscience 2, 349–354. doi:10.1038/ngeo475.

Kelley, C. A., Nicholson, B. E., Beaudoin, C. S., Detweiler, A. M., and Bebout, B. M. (2014). Trimethylamine and Organic Matter Additions Reverse Substrate Limitation Effects on the κ13C Values of Methane Produced in Hypersaline Microbial Mats. Appl. Environ. Microbiol. 80, 7316–7323. doi:10.1128/AEM.02641-14.

Kim, S.-J., Park, S.-J., Jung, M.-Y., Kim, J.-G., Min, U.-G., Hong, H.-J., et al. (2014). Draft genome sequence of an aromatic compound-degrading bacterium, Desulfobacula sp. TS, belonging to the Deltaproteobacteria. FEMS Microbiol Lett 360, 9–12. doi:10.1111/1574-6968.12591.

King, G. M. (1984a). Metabolism of Trimethylamine, Choline, and Glycine Betaine by Sulfate-Reducing and Methanogenic Bacteria in Marine Sediments. Appl Environ Microbiol 48, 719– 725.

King, G. M. (1984b). Utilization of hydrogen, acetate, and “noncompetitive”; substrates by methanogenic bacteria in marine sediments. Geomicrobiology Journal 3, 275–306.

Kivenson, V., and Giovannoni, S. J. (2020). An Expanded Genetic Code Enables Trimethylamine Metabolism in Human Gut Bacteria. mSystems 5. doi:10.1128/mSystems.00413-20.

Kivenson, V., Lemkau, K. L., Pizarro, O., Yoerger, D. R., Kaiser, C., Nelson, R. K., et al. (2019). Ocean Dumping of Containerized DDT Waste Was a Sloppy Process. Environ. Sci. Technol. doi:10.1021/acs.est.8b05859.

Kozich, J. J., Westcott, S. L., Baxter, N. T., Highlander, S. K., and Schloss, P. D. (2013). Development of a Dual-Index Sequencing Strategy and Curation Pipeline for Analyzing Amplicon Sequence Data on the MiSeq Illumina Sequencing Platform. Appl Environ Microbiol 79, 5112–5120. doi:10.1128/AEM.01043-13.

Kozlov, A. M., Darriba, D., Flouri, T., Morel, B., and Stamatakis, A. (2019). RAxML-NG: a fast, scalable and user-friendly tool for maximum likelihood phylogenetic inference. Bioinformatics 35, 4453–4455.

Krzycki, J. A. (2013). The path of lysine to pyrrolysine. Curr Opin Chem Biol 17, 619–625. doi:10.1016/j.cbpa.2013.06.023.

Kuever, J., Könneke, M., Galushko, A., and Drzyzga, O. (2001). Reclassification of Desulfobacterium phenolicum as Desulfobacula phenolica comb. nov. and description of strain SaxT as Desulfotignum balticum gen. nov., sp. nov. International Journal of Systematic and Evolutionary Microbiology 51, 171–177. doi:10.1099/00207713-51-1-171.

Lagesen, K., Hallin, P., Rødland, E. A., Stærfeldt, H.-H., Rognes, T., and Ussery, D. W. (2007). RNAmmer: consistent and rapid annotation of ribosomal RNA genes. Nucleic acids research 35, 3100–3108.

Langmead, B., Trapnell, C., Pop, M., and Salzberg, S. L. (2009). Ultrafast and memory-efficient alignment of short DNA sequences to the human genome. Genome Biology 10, R25. doi:10.1186/gb-2009-10-3-r25.

Laslett, D., and Canback, B. (2004). ARAGORN, a program to detect tRNA genes and tmRNA genes in nucleotide sequences. Nucleic Acids Res 32, 11–16. doi:10.1093/nar/gkh152.

Li, D., Liu, C.-M., Luo, R., Sadakane, K., and Lam, T.-W. (2015). MEGAHIT: an ultra-fast single-node solution for large and complex metagenomics assembly via succinct de Bruijn graph. Bioinformatics 31, 1674–1676. doi:10.1093/bioinformatics/btv033.

Li, H., Handsaker, B., Wysoker, A., Fennell, T., Ruan, J., Homer, N., et al. (2009). The Sequence Alignment/Map format and SAMtools. Bioinformatics 25, 2078–2079. doi:10.1093/bioinformatics/btp352.

Li, W., and Godzik, A. (2006). Cd-hit: a fast program for clustering and comparing large sets of protein or nucleotide sequences. Bioinformatics 22, 1658–1659.

Liu, Z., Reches, M., Groisman, I., and Engelberg-Kulka, H. (1998). The nature of the minimal ‘selenocysteine insertion sequence’(SECIS) in Escherichia coli. Nucleic acids research 26, 896–902.

Lovley, D. R., and Klug, M. J. (1983). Methanogenesis from methanol and methylamines and acetogenesis from hydrogen and carbon dioxide in the sediments of a eutrophic lake. Applied and Environmental Microbiology 45, 1310–1315.

Low, S. C., and Berry, M. J. (1996). Knowing when not to stop: selenocysteine incorporation in eukaryotes. Trends in biochemical sciences 21, 203–208.

McMurdie, P. J., and Holmes, S. (2013). phyloseq: an R package for reproducible interactive analysis and graphics of microbiome census data. PLoS ONE 8, e61217. doi:10.1371/journal.pone.0061217.

Monteverde, D., Sylvan, J., Suffridge, C., Baronas, J. J., Fichot, E., Fuhrman, J., et al. (2018). Distribution of extracellular flavins in a coastal marine basin and their relationship to redox gradients and microbial community members. Environ. Sci. Technol. doi:10.1021/acs.est.8b02822.

Mukai, T., Englert, M., Tripp, H. J., Miller, C., Ivanova, N. N., Rubin, E. M., et al. (2016). Facile Recoding of Selenocysteine in Nature. Angewandte Chemie International Edition 55, 5337– 5341. doi:10.1002/anie.201511657.

Mukherjee, S., Stamatis, D., Bertsch, J., Ovchinnikova, G., Sundaramurthi, J. C., Lee, J., et al. (2021). Genomes OnLine Database (GOLD) v.8: overview and updates. Nucleic Acids Research 49, D723–D733. doi:10.1093/nar/gkaa983.

Namy, O., Zhou, Y., Gundllapalli, S., Polycarpo, C. R., Denise, A., Rousset, J.-P., et al. (2007). Adding pyrrolysine to the Escherichia coli genetic code. FEBS letters 581, 5282–5288.

Nawrocki, E. P., and Eddy, S. R. (2013). Infernal 1.1: 100-fold faster RNA homology searches. Bioinformatics 29, 2933–2935. doi:10.1093/bioinformatics/btt509.

Nystrom, N. A., Levine, M. J., Roskies, R. Z., and Scott, J. (2015). Bridges: a uniquely flexible HPC resource for new communities and data analytics. in Proceedings of the 2015 XSEDE Conference: Scientific Advancements Enabled by Enhanced Cyberinfrastructure (ACM), 30.

Orcutt, B., Sylvan, J., Knab, N., and Edwards, K. (2011). Microbial ecology of the dark ocean above, at, and below the seafloor. Microbiol Mol Biol Rev 75, 361–422.

Orsi, W. D., Edgcomb, V. P., Christman, G. D., and Biddle, J. F. (2013). Gene expression in the deep biosphere. Nature 499, 205–208. doi:10.1038/nature12230.

Paul, L., Ferguson, D. J., and Krzycki, J. A. (2000). The trimethylamine methyltransferase gene and multiple dimethylamine methyltransferase genes of Methanosarcina barkeri contain in-frame and read-through amber codons. Journal of bacteriology 182, 2520–2529.

Paul, L., and Krzycki, J. A. (1996). Sequence and transcript analysis of a novel Methanosarcina barkeri methyltransferase II homolog and its associated corrinoid protein homologous to methionine synthase. J. Bacteriol. 178, 6599–6607. doi:10.1128/jb.178.22.6599-6607.1996.

Peng, T., Lin, J., Xu, Y.-Z., and Zhang, Y. (2016). Comparative genomics reveals new evolutionary and ecological patterns of selenium utilization in bacteria. The ISME Journal 10, 2048–2059. doi:10.1038/ismej.2015.246.

Polycarpo, C., Ambrogelly, A., Bérubé, A., Winbush, S. M., McCloskey, J. A., Crain, P. F., et al. (2004). An aminoacyl-tRNA synthetase that specifically activates pyrrolysine. PNAS 101, 12450–12454. doi:10.1073/pnas.0405362101.

Porter, A. W., and Young, L. Y. (2013). The bamA gene for anaerobic ring fission is widely distributed in the environment. Frontiers in microbiology 4, 302.

Prat, L., Heinemann, I. U., Aerni, H. R., Rinehart, J., O’Donoghue, P., and Söll, D. (2012). Carbon source-dependent expansion of the genetic code in bacteria. PNAS 109, 21070–21075. doi:10.1073/pnas.1218613110.

Price, M. N., Dehal, P. S., and Arkin, A. P. (2010). FastTree 2–approximately maximum-likelihood trees for large alignments. PloS one 5, e9490.

Pruesse, E., Quast, C., Knittel, K., Fuchs, B. M., Ludwig, W., Peplies, J., et al. (2007). SILVA: a comprehensive online resource for quality checked and aligned ribosomal RNA sequence data compatible with ARB. Nucleic acids research 35, 7188–7196.

Reeburgh, W. S. (2007). Oceanic Methane Biogeochemistry. Chem. Rev. 107, 486–513. doi:10.1021/cr050362v.

Rho, M., Tang, H., and Ye, Y. (2010). FragGeneScan: predicting genes in short and error-prone reads. Nucleic Acids Res. 38, e191. doi:10.1093/nar/gkq747.

Rodriguez-R, L. M., and Konstantinidis, K. T. (2014). Bypassing cultivation to identify bacterial species. Microbe 9, 111–118.

Rodriguez-R, L. M., and Konstantinidis, K. T. (2016). The enveomics collection: a toolbox for specialized analyses of microbial genomes and metagenomes. PeerJ Inc. doi:10.7287/peerj.preprints.1900v1.

Santesmasses, D., Mariotti, M., and Guigó, R. (2017). Computational identification of the selenocysteine tRNA (tRNASec) in genomes. PLOS Computational Biology 13, e1005383. doi:10.1371/journal.pcbi.1005383.

Sogin, M. L., Morrison, H. G., Huber, J. A., Welch, D. M., Huse, S. M., Neal, P. R., et al. (2006). Microbial diversity in the deep sea and the underexplored “rare biosphere.” Proceedings of the National Academy of Sciences 103, 12115–12120.

Srinivasan, G., James, C. M., and Krzycki, J. A. (2002a). Pyrrolysine Encoded by UAG in Archaea: Charging of a UAG-Decoding Specialized tRNA. Science 296, 1459–1462. doi:10.1126/science.1069588.

Srinivasan, G., James, C. M., and Krzycki, J. A. (2002b). Pyrrolysine Encoded by UAG in Archaea: Charging of a UAG-Decoding Specialized tRNA. Science 296, 1459–1462. doi:10.1126/science.1069588.

Stadtman, T. C. (1996). Selenocysteine. Annual review of biochemistry 65, 83–100.

Sun, J., Mausz, M. A., Chen, Y., and Giovannoni, S. J. (2019). Microbial trimethylamine metabolism in marine environments. Environmental Microbiology 21, 513–520. doi:10.1111/1462-2920.14461.

Thureborn, P., Franzetti, A., Lundin, D., and Sjöling, S. (2016). Reconstructing ecosystem functions of the active microbial community of the Baltic Sea oxygen depleted sediments. PeerJ 4, e1593. doi:10.7717/peerj.1593.

Ticak, T., Kountz, D. J., Girosky, K. E., Krzycki, J. A., and Ferguson, D. J. (2014). A nonpyrrolysine member of the widely distributed trimethylamine methyltransferase family is a glycine betaine methyltransferase. Proceedings of the National Academy of Sciences 111, E4668– E4676.

Towns, J., Cockerill, T., Dahan, M., Foster, I., Gaither, K., Grimshaw, A., et al. (2014). XSEDE: accelerating scientific discovery. Computing in Science & Engineering 16, 62–74.

Valentine, D. L., Fisher, G. B., Pizarro, O., Kaiser, C. L., Yoerger, D., Breier, J. A., et al. (2016). Autonomous Marine Robotic Technology Reveals an Expansive Benthic Bacterial Community Relevant to Regional Nitrogen Biogeochemistry. Environ. Sci. Technol. 50, 11057–11065. doi:10.1021/acs.est.6b03584.

Vannelli, T., Messmer, M., Studer, A., Vuilleumier, S., and Leisinger, T. (1999). A corrinoid-dependent catabolic pathway for growth of a Methylobacterium strain with chloromethane. Proc Natl Acad Sci U S A 96, 4615–4620.

Wöhlbrand, L., Jacob, J. H., Kube, M., Mussmann, M., Jarling, R., Beck, A., et al. Complete genome, catabolic sub-proteomes and key-metabolites of Desulfobacula toluolica Tol2, a marine, aromatic compound-degrading, sulfate-reducing bacterium. Environmental Microbiology 15, 1334–1355. doi:10.1111/j.1462-2920.2012.02885.x.

Xiao, K.-Q., Beulig, F., Røy, H., Jørgensen, B. B., and Risgaard-Petersen, N. (2018). Methylotrophic methanogenesis fuels cryptic methane cycling in marine surface sediment. Limnology and Oceanography 63, 1519–1527. doi:10.1002/lno.10788.

Zhang, Y., Fomenko, D. E., and Gladyshev, V. N. (2005). The microbial selenoproteome of the Sargasso Sea. Genome Biol 6, R37. doi:10.1186/gb-2005-6-4-r37.

Zhang, Y., and Gladyshev, V. N. (2005). An algorithm for identification of bacterial selenocysteine insertion sequence elements and selenoprotein genes. Bioinformatics 21, 2580–2589. doi:10.1093/bioinformatics/bti400.

Zhang, Y., and Gladyshev, V. N. (2007). High content of proteins containing 21st and 22nd amino acids, selenocysteine and pyrrolysine, in a symbiotic deltaproteobacterium of gutless worm Olavius algarvensis. Nucleic acids research 35, 4952–4963.

Zhang, Y., and Gladyshev, V. N. (2008). Trends in Selenium Utilization in Marine Microbial World Revealed through the Analysis of the Global Ocean Sampling (GOS) Project. PLOS Genetics 4, e1000095. doi:10.1371/journal.pgen.1000095.

Zhuang, G.-C., Elling, F. J., Nigro, L. M., Samarkin, V., Joye, S. B., Teske, A., et al. (2016). Multiple evidence for methylotrophic methanogenesis as the dominant methanogenic pathway in hypersaline sediments from the Orca Basin, Gulf of Mexico. Geochimica et Cosmochimica Acta 187, 1–20. doi:10.1016/j.gca.2016.05.005.

Zinoni, F., Birkmann, A., Leinfelder, W., and Böck, A. (1987). Cotranslational insertion of selenocysteine into formate dehydrogenase from Escherichia coli directed by a UGA codon. PNAS 84, 3156–3160. doi:10.1073/pnas.84.10.3156.

Zinoni, F., Heider, J., and Böck, A. (1990). Features of the formate dehydrogenase mRNA necessary for decoding of the UGA codon as selenocysteine. Proceedings of the National Academy of Sciences 87, 4660–4664.

