## Supplemental Information for "An ecological basis for dual genetic code expansion in marine deltaproteobacteria"

1) Figures S1-S6
2) Tables S1-S10

**SI Figures**

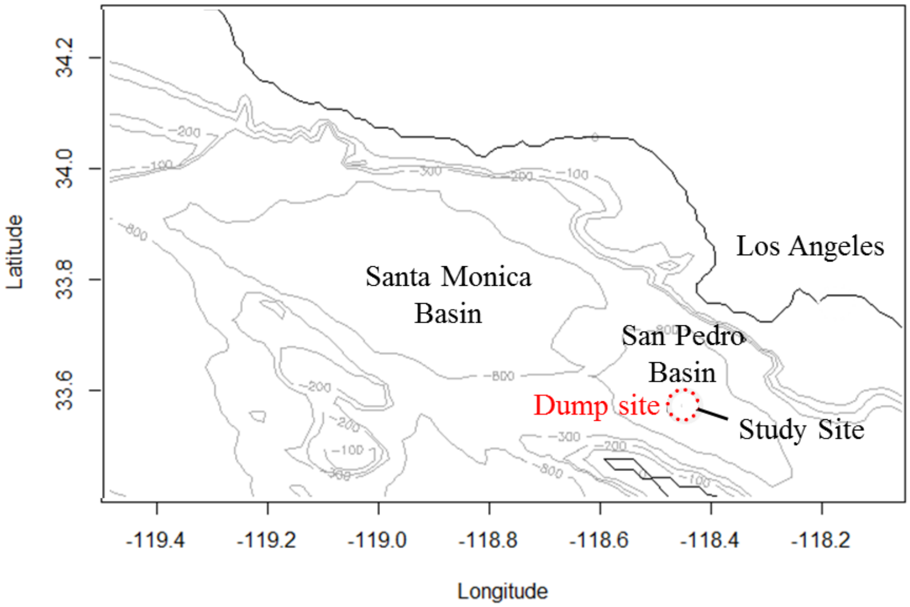


**Figure S1)** Location of the barrel dumpsite and the study site off of the coast of California.

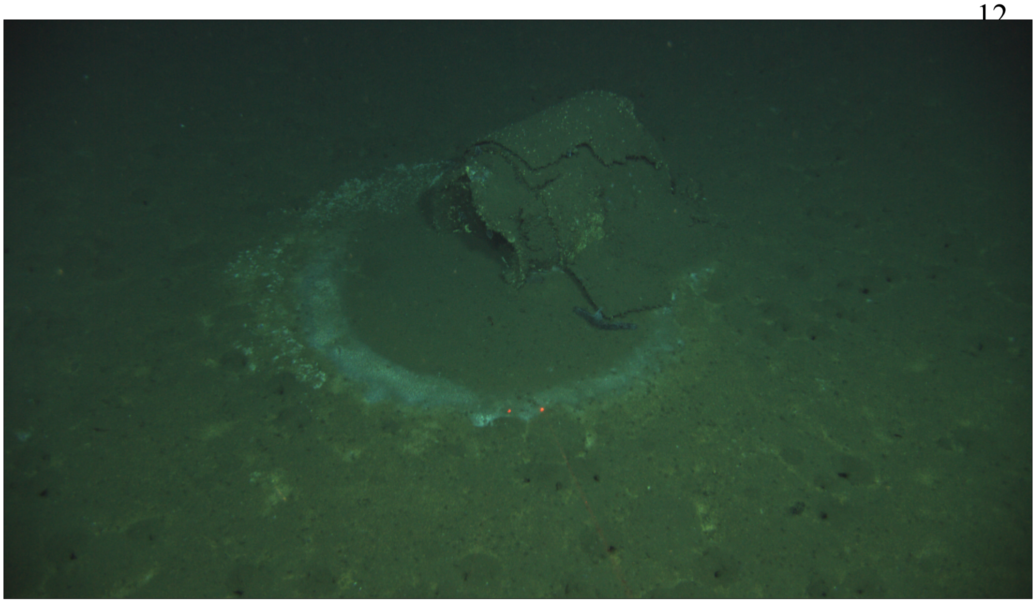

**Figure S2)** Apparent infauna burrows visible in the sediment near a barrel, and absent between the barrel and the microbial mat ring.

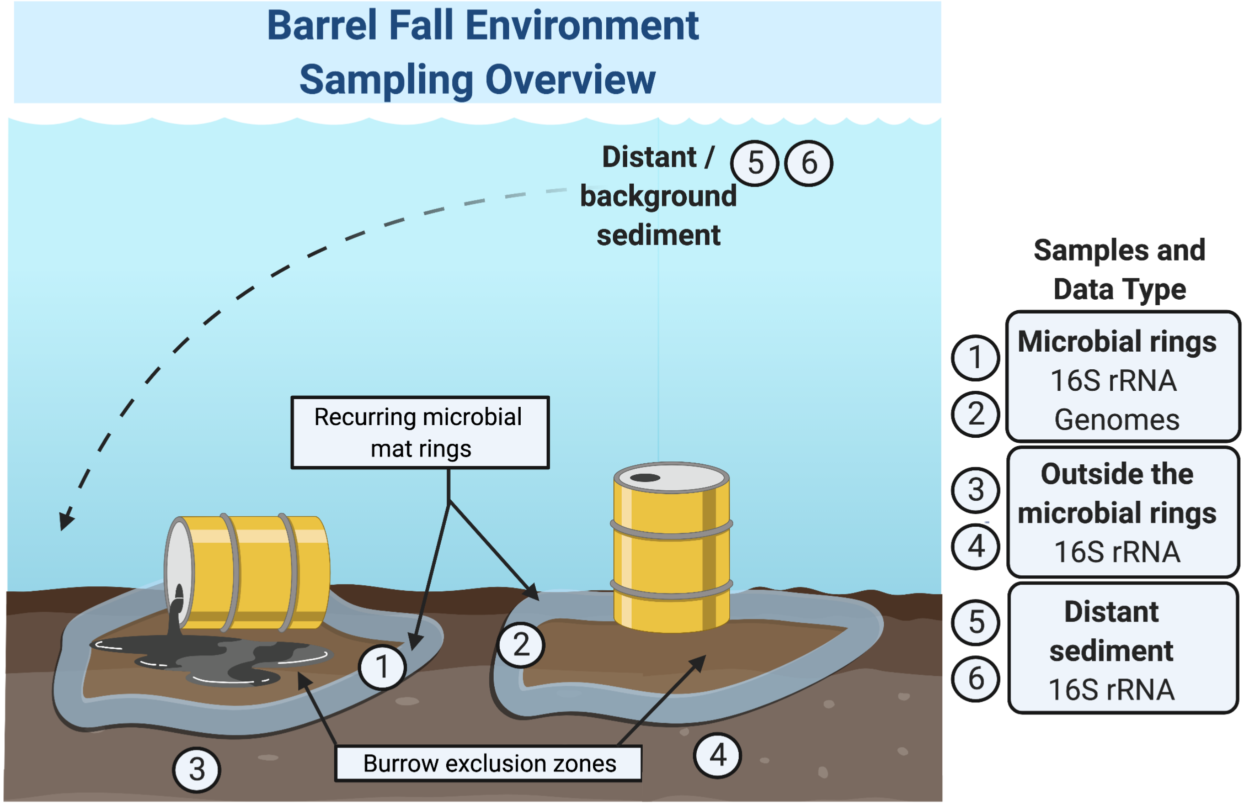


**Figure S3)** Sampling schematic describing the site and sequencing methods used for microbial analyses.

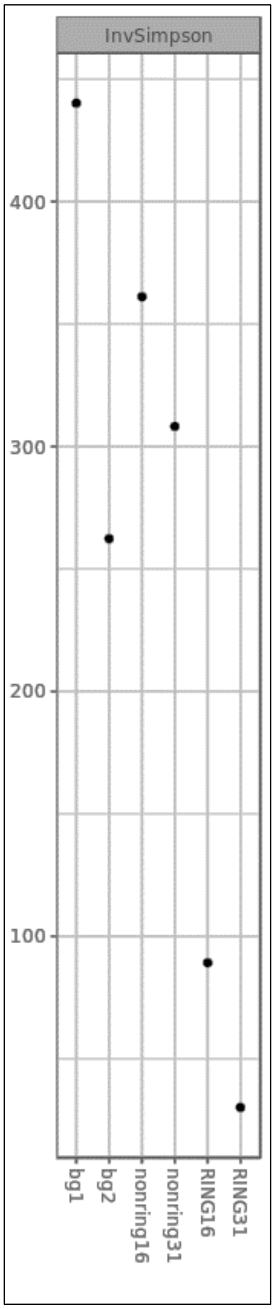


**Figure S4)** Inverse Simpsons index for diversity (bg1, bg2: background 1 and 2; nonring 16 and 31: outside of the microbial ring at bbl 16 and bbl 31; RING16: microbial mat at bbl 16; RING31: microbial mat at bbl 31).

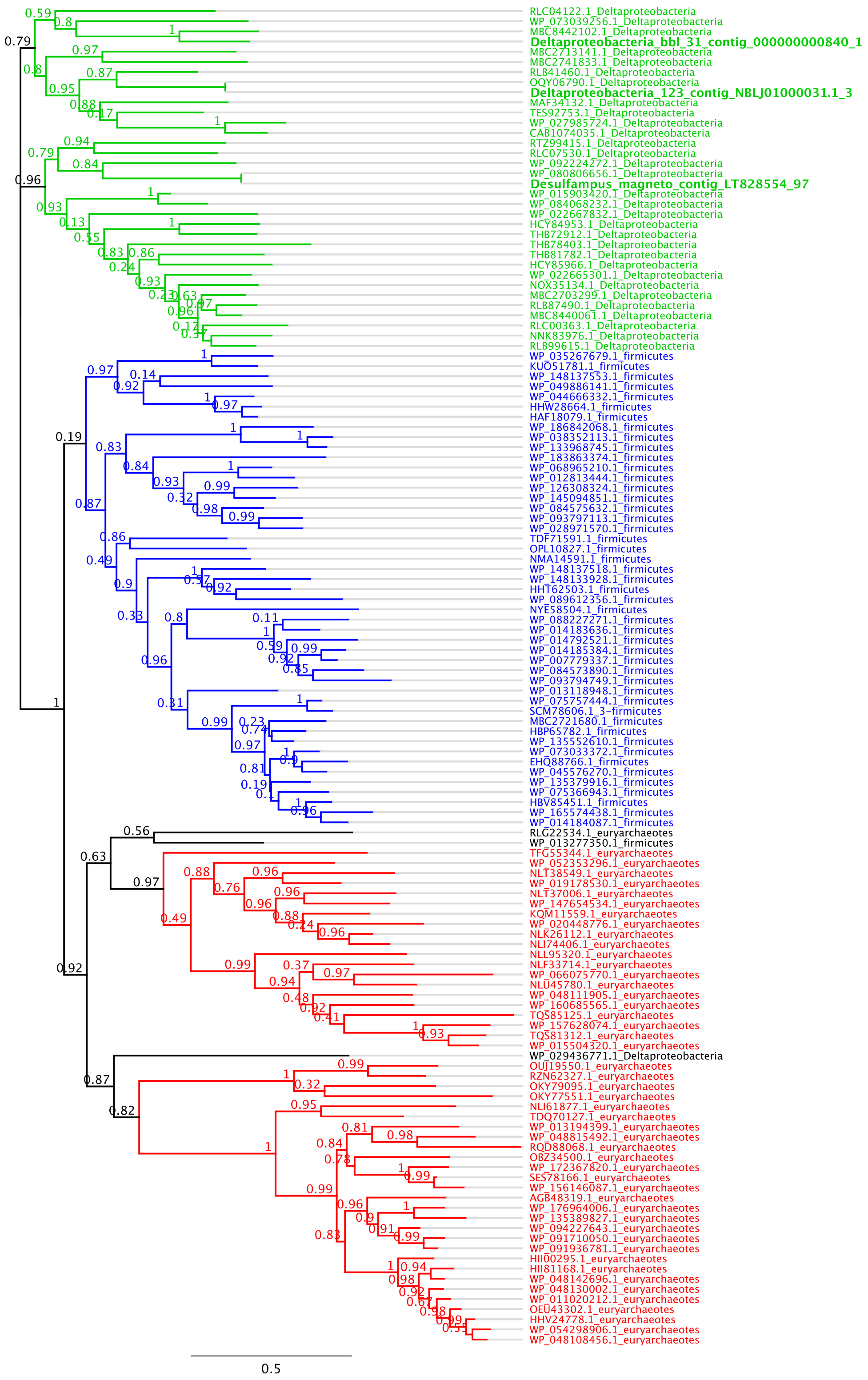


**Fig S5)** Phylogeny of PylB reconstructed using FastTree. This tree shows a horizontal view of the tree from **Fig. 3A**. Branch support values are shown at all nodes across the tree. Major clades are highlighted according to taxonomic classification, in either red (Euryarchaeota), blue (Firmicutes), or green (Deltaproteobacteria).

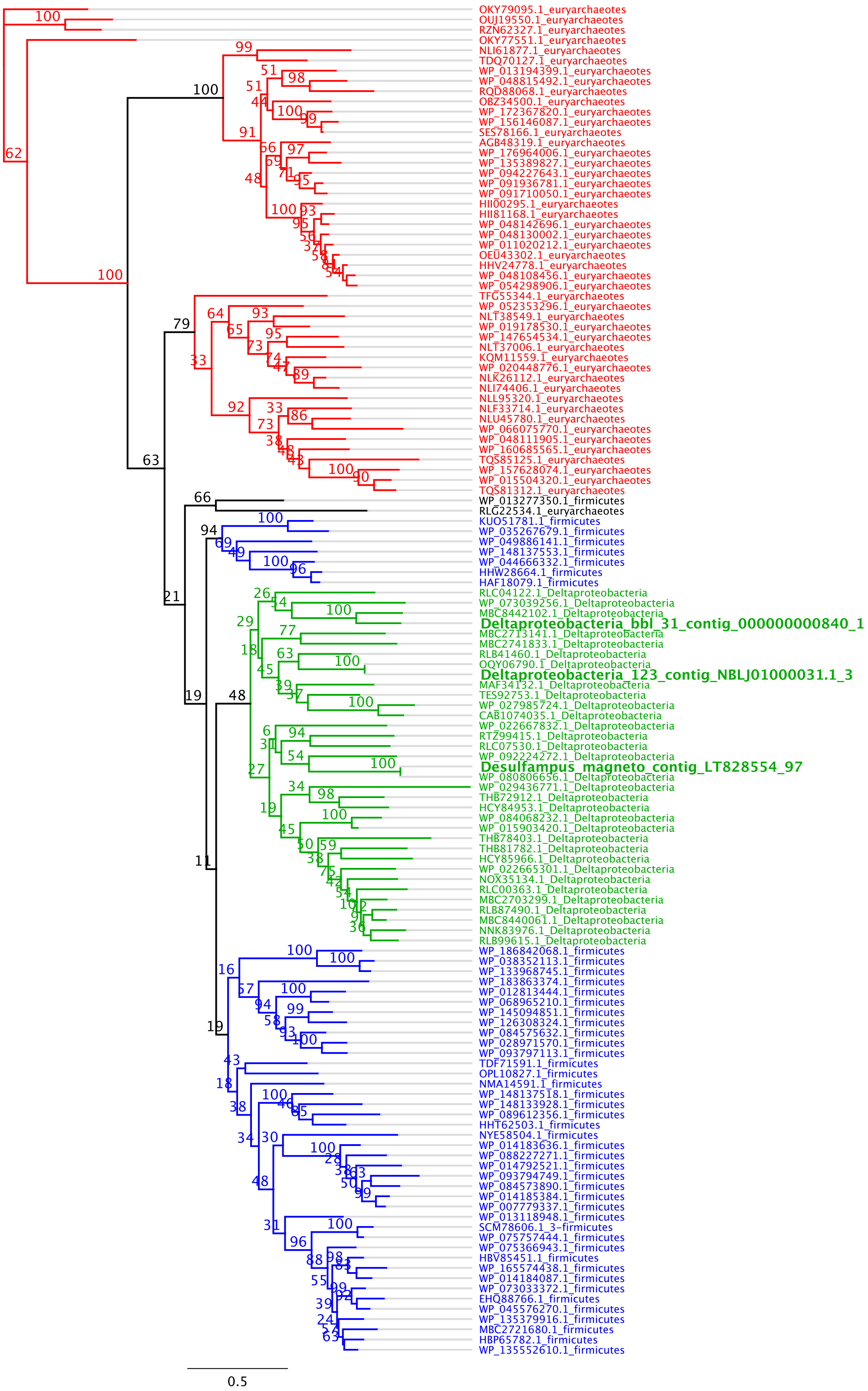


**Fig S6)** Phylogeny of PylB reconstructed using RaxML. Branch support values are shown at all nodes across the tree. Major clades are highlighted according to taxonomic classification, in either red (Euryarchaeota), blue (Firmicutes), or green (Deltaproteobacteria).

### SI Tables

**Table S1.** Percent abundance of top twenty taxa by location with ASV number as indicated.

| ASV | bg1 | bg2 | Nonring bbl-16 | Nonring bbl-31 | Ring bbl-16 | Ring bbl- 31 |
|---|---|---|---|---|---|---|
| 1 | 0 | 0 | 0 | 0 | 8 | 16 |
| 2 | 3 | 0 | 3 | 2 | 0 | 0 |
| 3 | 1 | 3 | 0 | 1 | 0 | 2 |
| 4 | 0 | 0 | 0 | 0 | 1 | 4 |
| 5 | 0 | 0 | 1 | 0 | 4 | 1 |
| 6 | 0 | 0 | 0 | 0 | 0 | 4 |
| 7 | 0 | 1 | 0 | 0 | 2 | 2 |
| 8 | 1 | 0 | 2 | 2 | 0 | 0 |
| 9 | 0 | 0 | 0 | 0 | 2 | 2 |
| 10 | 1 | 1 | 1 | 1 | 0 | 1 |
| 11 | 1 | 1 | 1 | 1 | 1 | 0 |
| 12 | 0 | 0 | 0 | 0 | 1 | 1 |
| 13 | 0 | 0 | 0 | 0 | 1 | 2 |
| 14 | 1 | 1 | 1 | 1 | 0 | 0 |
| 15 | 1 | 1 | 0 | 1 | 0 | 0 |
| 16 | 0 | 1 | 0 | 1 | 0 | 1 |
| 17 | 0 | 0 | 0 | 0 | 2 | 1 |
| 18 | 0 | 0 | 0 | 0 | 0 | 2 |
| 19 | 0 | 1 | 0 | 0 | 0 | 0 |
| 20 | 1 | 0 | 1 | 0 | 0 | 0 |

**Table S1 continued.** Matching taxonomy for each ASV of the top twenty taxa. The phylum, class, family, and genus are shown. NA indicates unclassified at the given taxonomic level.

| ASV | Phylum | Class | Family | Genus |
|---|---|---|---|---|
| 1 | Proteobacteria | Deltaproteobacteria | Desulfobacteraceae | Desulfobacula |
| 2 | Proteobacteria | Gammaproteobacteria | NA | NA |
| 3 | Lokiarchaeota | NA | NA | NA |
| 4 | Latescibacteria | NA | NA | NA |
| 5 | Planctomycetes | Phycisphaerae | NA | NA |
| 6 | Bacteroidetes | Bacteroidia | Marinilabiaceae | NA |
| 7 | Lokiarchaeota | NA | NA | NA |
| 8 | Planctomycetes | Planctomycetacia | Brocadiaceae | Candidatus_Scalindua |
| 9 | Acetothermia | NA | NA | NA |
| 10 | Lokiarchaeota | NA | NA | NA |
| 11 | Lokiarchaeota | NA | NA | NA |
| 12 | Chloroflexi | Anaerolineae | Anaerolineaceae | NA |
| 13 | Spirochaetae | Spirochaetes | Spirochaetaceae | Spirochaeta_2 |
| 14 | Proteobacteria | Deltaproteobacteria | Desulfobacteraceae | Sva0081_sediment_group |
| 15 | Bacteroidetes | Bacteroidetes_BD2-2 | NA | NA |
| 16 | Chloroflexi | Anaerolineae | Anaerolineaceae | NA |
| 17 | Chloroflexi | Anaerolineae | Anaerolineaceae | NA |
| 18 | Lokiarchaeota | NA | NA | NA |
| 19 | Chloroflexi | Anaerolineae | Anaerolineaceae | NA |
| 20 | Chloroflexi | Anaerolineae | Anaerolineaceae | NA |

**Table S1 continued.** The corresponding 16S rRNA sequence for each ASV.

| **ASV #** | **Corresponding 16S rRNA nucleotide sequence** |
|---|---|
| 1 | CACGGGGGGCGCAAGCGTTATTCGGAATTATTGGGCGTAAAGGGCGCGTAGGCGGTCTTGTCGGTCAGATGTGAAAGCCCAGGGCTCAACCCTGGACGTGCATTTGAAACAGCAAGACTTGAGTACGGGAGAGGAAAGCGGAATTCCTGGTGTAGAGGTGAAATTCGTAGATATCAGGAGGAACACCGATGGCGAAGGCAGCTTTCTGGACCGATACTGACGCTGAGGCGCGAAGGCGTGGGTAGCGAACAGG |
| 2 | TACGGAGGGTGCAAGCGTTAATCGGAATTACTGGGCGTAAAGCGCTCGTAGGCGGTTTGTTAAGTCGGATGTGAAAGCCCCGGGCTCAACCTGGGAACTGCATTCGATACTGGCAAACTAGAGTATAGAAGAGGCAAGTGGAATTCCGGGTGTAGCGGTGAAATGCGTAGATATCCGGAGGAACATCAGTGGCGAAGGCGACTTGCTGGTCTAATACTGACGCTGAGGAGCGAAAGCGTGGGGAGCAAACGGG |
| 3 | AACCAGCTCTTCAAGTGGTCGGGATAATTATTGGGCTTAAAGTGTCCGTAGCCGGTTTAGTAAGTTCCTGGTAAAATCGGGTAGCTTAACTATCTATATGCTAGGAATACTACTATACTAGAGGGCGGGAGAGGTCTGAGGTACTACAGGGGTAGGGGTGAAATCTTATAATCCTTGTTGGACCACCAGTGGCGAAGGCGTCAGACTGGAACGCGCCTGACGGTGAGGGACGAAAGCCAGGGGAGCGAACCGG |
| 4 | TACGGAGGGTGCAAGCGTTGTTCGGATTTACTGGGTATAAAGGGTGCGCAGGCGGCCTGATAAGTCAGGGGTGAAATATGACGGCTCAACCGTCAAACTGCCCCTGAAACTGCCAGGCTTGAGTCCGAGAGAGGTAGGTGGAATTCCAGGTGTAGCGGTGAAATGCGTAAATATCTGGAGGAACACCGGTGGCGAAGGCGGCCTACTGGCTCGGAACTGACGCTCAGGCACGAAAGCTAGGGGAGCGAACGGG |
| 5 | TACGAAGGTGGCAAGCGTTGTTCGGAATCACTGGGCTTAAAGCGCACGCAGGCGGAAAAGAAAGTGTGGAGTGAAATCCCTCGGCTTAACCGGGGAACTGCTCTGCAAACTACTTTTCTTGAGGCAAGTAGGGGTACATGGAACTCTTGGTGGAGCGGTGGAATGCGTAGATATCAAGAGGAACGCCGATGGTGAAGACAGTGTACTGGGCTTGTCCTGACGCTGAGGTGCGAAAGCGTGGGGAGCGAACGGG |
| 6 | TACGGAGGGTGCGAGCGTTATCCGGATTTATTGGGTTTAAAGGGTGCGTAGGCGGAATATTAAGTCAGTGGTGAAATCCTGTGGCTCAACCATAGAATTGCCATTGATACTGATATTCTTGAATGCAGTTGAGGCAGGCGGAATGTGTAATGTAGCGGTGAAATGCTTAGATATTACACAGAACACCGATTGCGAAGGCAGCTTGCTAAACTGTGATTGACGCTGATGCACGAAAGCGTGGGGAGCGAACAGG |
| 7 | AACCAGCTCTTCAAGTGGTCGGGAATATTATTGGGCTTAAAGTGTCCGTAGCCGGTTTGGTAAGTTCCTGGTTAAATCTGGCAGCTTAACTGTCAGTCAGCTAGGAATACTACTTTACTAGAGGGTGGGAAAGGTTTGAGGTACTCCAGGGGTAGCGGTGAAATGCGATAATCCTTGGGGGACCACCAGTGGCGAAGGCGTCAGACTGGAACACGCCTGACGGTGAGGGACGAAAGCCAGGGGAGCGAACGGG |

**Table S1 continued.** The corresponding 16S rRNA sequence for each ASV.

| ASV # | Corresponding 16S rRNA nucleotide sequence |
|---|---|
| 8 | TACAGAGGTGGCAAGCGTTGTTCGGAATTATTGGGCGTAAAGAGCACGTAGGTGGGTTTGTAAGTCAGATGTGAAAGCCTTCTGTTCAACGGAAGAATTGCATCTGAAACTGCGAGTCTTGAGTGTAGGAGGGGAGAATGGAACTTCTGGTGGAGCGGTGAAATGCGTAGATATCAGAAGGAACGCCGGCGGCGAAAGCGATTCTCTGGCCTATTACTGACACTCAGTGTGCGAAAGCTAGGGGAGCAAACGGG |
| 9 | GACGAGGGATGCAAGCGTTATCCGGAATTACTGGGCGTAAAGGACGTCTAGGCGGTTGGATAAGTCATTTGTGAAATCCCAGGGCTTAACCCTGGAAGGTCTTGTGATACTGTCCGGCTTGGGTGTAGGAGAGGAGAGCGGAACTCACAGAGTAGCGGTGGAATGCGTAGATACTGTGAGGTACCCCGATGGCGAAGGCAGCTCTCTGGCCTATTACCGACGCTGAAGCGTGAAAGCGTGGGGAGCAAAGGGG |
| 10 | AACCAGCTCTTCAAGTGGTCGGGATTATTATTGGGCTTAAAGTGTTCGTAGCCTGTTTAGTAAGTTCTTGGTTAAATCGGATAGCTTAACTATCTGTCTGCTAAGAATACTACTATACTAGGGGGCGGGAGAGGTCTGAGGTACTCCAGGGGTAGCGGTGAAATGCTATAATCCTTGGGGGACCACCAGTGGCGAAGGCGTCAGACTGGAACGCGCCCGACGGTGAGGGACGAAAGCCAGGGGAGCGAACCGG |
| 11 | AACCAGCTCTTCAAGTGGTCGGGATTATTATTGGGCTTAAAGTGTTCGTAGCCTGTTTAGTAAGTTCCTGGTTAAATCGGATAGCTTAACTATCTGTCTGCTAGGAATACTACTATACTAGGGGGCGGGAGAGGTCTGAGGTACTCCAGGGGTAGCGGTGAAATGCTATAATCCTTGGGGGACCACCAGTGGCGAAGGCGTCAGACTGGAACGCGCCCGACGGTGAGGGACGAAAGCCAGGGGAGCGAACCGG |
| 12 | TACGTAGGAGGCGAGCGTTATCCGGATTTATTGGGCGTAAAGCGCGTGCAGGTGGTTTGGTAAGTTGGGTATGAAATCTTCTGGCTTAACTAGGAGAGGTTGCTCAAAACTGCCAGACTAGAGGACGATAGAGGAAGGTGGAATTCCCGGTGTAGTAGTGAAATGCGTAGATATCGGGAGGAACACCAGTGGCGAAGGCGGCCTTCTGGGTCGTTCCTGACACTAAGACGCGAAAGCATGGGTAGCAAACGGG |
| 13 | CACGTATGGGGCGAGCGTTGTTCGGAATCATTGGGCGTAAAGGGCGCGCAGGCGGTTATATAAGCCTGGTGTGAAATACTGCAGCTCAACTGCAGAACCGCACTGGGAACTGTATGACTGGAGTTCAAGAGGGGAAGCTGGAATTCCTGGTGTAGGGGTGAAATCTGTAGATATCAGGAAGAACATCAGTGGCGAAGGCGAGCTTCTGGCTATGAACTGACGCTGAGGCGCGAAAGCGTGGGGAGCAAACAGG |

**Table S1 continued.** The corresponding 16S rRNA sequence for each ASV.

| ASV # | Corresponding 16S rRNA nucleotide sequence |
|---|---|
| 14 | CACGGGGGGTGCAAGCGTTATTCGGAATCACTGGGCGTAAAGAGCGCGTAGGCGGTCTCTTAAGTCAGATGTGAAAGCCCGGGGCTCAACCCCGGAAGTGCATTTGAAACGAAGGGACTTGAGTATGGGAGAGGGAAGTGGAATTCCTGGTGTAGCGGTGAAATGCGTAGATATCAGGAGGAACACCGGTGGCGATGGCGACTTCCTGGACCAATACTGACGCTGAGGCGCGAAGGCGTGGGGAGCAAACAGG |
| 15 | TACGGAGGATGCAAGCGTTATCCGGATTTATTGGGTTTAAAGGGTACGTAGGCGGAAAATTAAGTCAGTAGTGAAATCCTGCAGCTTAACTGTAGAACTGTTATTGATACTGGTTTTCTTGAATATAGTTGAGGTAGGCGGAATGTGTAATGTAGCGGTGAAATGCTTAGATATTACACAGAACACCGATTGCGAAGGCAGCTTACTAAGCTATGATTGACGCTGAGGTACGAAAGCGTGGGGAGCGAACAGG |
| 16 | AACGTAGGATCCGAGCGTTATCCGAATTCACTGGGCGTAAAGCGCGTGTAGGCGGTTCGGTAAGTTGGATGTGAAAGCTCCCGGCTCAACTGGGAGAGGACGTTCAAAACTGTTGGACTAGAGGGCGGAAGAGGGAGGTGGAATTCCCGGTGTAGTGGTGAAATGCGTAGATATCGGGAGGAACACCAGTGGCGAAGGCGGCCTCCTGGGCCGCACCTGACGCTCAGACGCGAAAGCTAGGGTAGCAAACGGG |
| 17 | TACGTAGGAGGCAAGCGTTATCCGGATTCATTGGGCGTAAAGCGCGTGCAGGTGGTTTGGTAAGTTGGGTATGAAATCTTCTGGCTTAACTAGGAGAGGTTGCTCAAAACTGTCAGACTAGAGGACGATAGAGGAAGGTGGAATTCCCGGTGTAGTAGTGAAATGCGTAGATATCGGGAGGAACACCAGTGGCGAAGGCGGCCTTCTGGGTCGTTCCTGACACTAAGACGCGAAAGCATGGGTAGCAAACGGG |
| 18 | AACCAGCTCTTCAAGTGGTCGGGAATATTATTGGGCTTAAAGTGTCCGTAGCCGGTTTGAACAGTTCCTGGTTAAATCTGGTAGCTTAACTATCAGTCAGCTAGGAATACTATCTTACTAGAGGGTGGGAAAGGCTTGGGGTACTCCGGGGGTAGCGGTGAAATGCGATAATCCTCGGGGGACCACCAGTGGCGAAGGCGCCAAGCTGGAACACGCCTGACGGTGAGGGACGAAAGCCAGGGGAGCGAACGGG |
| 19 | CACGTAGGATCCGAGCGTTATCCGAATTTACTGGGCGTAAAGCGCGTGTAGGCGGCCGGGTAAGTTGGACGTGAAAGCTCCTGGCTCAACTAGGAGAGGTCGTTCAAAACTGCCTGGCTAGAGGGCGACAGAGGGAGGTGGAATTCCCGGTGTAGTGGTGAAATGCGTAGATATCGGGAGGAACACCAGTGGCGAAGGCGGCCTCCTGGGTCGCCCCTGACGCTCAGACGCGAAAGCTAGGGGAGCAAACGGG |
| 20 | GACATAGGAGGCGAGCGTTATCCGGATTTATTGGGCGTAAAGTGCGTTGAGGCGGCATTGTAAGTTGGACGTGAAAGCTCCCGGCTTAACTGGGAGAGGTCGTTCAATACTGCAAGGCTAGAGGGCAGTAGAGGGGGGTGGAATTCCCGGTGTAGTGGTGAAATGCGTAGATATCGGGAGGAACACCAGTGGCGAAGGCGGCCCCCTGGACTGTTACTGACGCTGAAGGCGAAAGCTAGGGTAGCAAACGGG |

**Table S2**. Assembly statistics following whole genome sequencing. Assembly shown was performed with Megahit with standard parameters unless otherwise indicated (with minimum contig length of 1000 bp).

| Sample name | Raw reads file size | Total assembly length | Max contig length | Mean contig length | N50 | Most abundant 16S ASV percent abundance | Reconstructed genome(s)? |
|---|---|---|---|---|---|---|---|
| Core 12 (background) | 10.6 G | 400,023 | 10,690 | 1,481 | 1,359 | 3% | No |
| Core 13 (background) | 9.2 G | 9,475,213 | 34,633 | 1,364 | 1,288 | 3% | No |
| Core 18 (outside ring, bbl-31) | 9.2 G | 27,849,371 | 16,830 | 1,575 | 1,506 | 3% | No |
| Core 7 (outside ring, bbl-16) | 12.6 G | 9,734,423 | 43,818 | 1,466 | 1,360 | 3% | No |
| Core 19 (ring, bbl-31) | 15.2 G | 106,625,211 | 167,842 | 2,185 | 2,228 | 16% | Yes |
| Core 8 (ring, bbl-16) | 22 G | 126,963,432 | 331,719 | 1,972 | 1,957 | 8% | Yes |

**Table S3**. Properties and taxonomic identification of the reconstructed genomes.

| Bin name | Size Mb | Mean Coverage | GC % | NCBI Taxonomy (Order unless otherwise specificed) | Gene used for NCBI Taxonomy |
|---|---|---|---|---|---|
| CORE_8_RING_Bin_00001 (Deltaproteobacteria-bbl-16 ) | 2.55 | 83 | 41 | Desulfobacterales | recA |
| CORE_8_RING_Bin_00002 | 3.28 | 23 | 50 | Candidate division Zixibacteria (Phylum) | recA |
| CORE_8_RING_Bin_00003 | 2.27 | 13 | 39 | Bacteroidales | Rp_S8 |
| CORE_8_RING_Bin_00004 | 0.61 | 21 | 28 | Candidatus Woesearchaeota (Phylum) | Rp_S8 |
| CORE_8_RING_Bin_00005 | 2.50 | 12 | 44 | Candidatus Marinimicrobia (Phylum) | recA |
| CORE_19_RING_Bin_ 00001 (Deltaproteobacteria-bbl-31) | 2.82 | 123 | 41 | Desulfobacterales | recA, 16S rRNA |
| CORE_19_RING_Bin_0000 2 | 3.39 | 45 | 32 | Bacteroidales | recA |
| CORE_19_RING_Bin_0000 3 | 3.78 | 15 | 51 | Candidatus Latescibacteria (Phylum) | recA |
| CORE_19_RING_Bin_0000 4 | 3.58 | 36 | 56 | Gemmatimonadales | recA |
| CORE_19_RING_Bin_0000 5 | 1.74 | 10 | 40 | Victivallales | recA |
| CORE_19_RING_Bin_0000 6 | 2.13 | 13 | 58 | Spirochaetales | recA |

**Table S4**. Taxonomic information from alternative database for reconstructed genomes.

| Bin name | GTDB-TK taxonomy (Rank as indicated) |
| --- | --- |
| CORE_8_RING_Bin_00001 (Deltaproteobacteria-bbl-16 ) | d__Bacteria;p__Desulfobacterota;c__Desulfobacteria; o__Desulfobacterales;f__Desulfobacteraceae;g__Desulfobacula |
| CORE_8_RING_Bin_00002 | d__Bacteria;p__Zixibacteria |
| CORE_8_RING_Bin_00003 | d__Bacteria;p__Bacteroidota;c__Bacteroidia;o__Bacteroidales |
| CORE_8_RING_Bin_00004 | d__Archaea;p__Nanoarchaeota;c__Nanoarchaeia;o__Woesearchaeales |
| CORE_8_RING_Bin_00005 | d__Bacteria;p__Marinisomatota |
| CORE_19_RING_Bin_00001 (Deltaproteobacteria-bbl-31) | d__Bacteria;p__Desulfobacterota;c__Desulfobacteria;o__ Desulfobacterales;f__Desulfobacteraceae;g__Desulfobacula |
| CORE_19_RING_Bin_00002 | d__Bacteria;p__Bacteroidota;c__Bacteroidia;o__Bacteroidales |
| CORE_19_RING_Bin_00003 | d__Bacteria;p__Krumholzibacteriota;c__Krumholzibacteria |
| CORE_19_RING_Bin_00004 | d__Bacteria;p__Krumholzibacteriota;c__Krumholzibacteria |
| CORE_19_RING_Bin_00005 | d__Bacteria;p__Verrucomicrobiota;c__Lentisphaeria;o__ Victivallales;f__Victivallaceae |
| CORE_19_RING_Bin_00006 | d__Bacteria;p__Spirochaetota;c__Spirochaetia;o__Spirochaetales;f__Alkalispirochaetaceae |

**Table S5.** Enumeration of COG categories for Deltaproteobacterial-bbl genomes.

| | *COG Category* | *Deltaproteo-bacteria -bbl-31* | *Deltaproteo-bacteria -bbl-16* |
|---|---|---|---|
| A | RNA processing and modification | 0 | 0 |
| B | Chromatin structure and dynamics | 3 | 3 |
| C | Energy production and conversion | 239 | 208 |
| D | Cell cycle control, cell division, chromosome partitioning | 24 | 23 |
| E | Amino acid transport and metabolism | 248 | 217 |
| F | Nucleotide transport and metabolism | 51 | 45 |
| G | Carbohydrate transport and metabolism | 89 | 84 |
| H | Coenzyme transport and metabolism | 108 | 94 |
| I | Lipid transport and metabolism | 93 | 84 |
| J | Translation, ribosomal structure and biogenesis | 156 | 148 |
| K | Transcription | 107 | 89 |
| L | Replication, recombination and repair | 91 | 75 |
| M | Cell wall/ membrane/ envelope biogenesis | 110 | 88 |
| N | Cell motility | 51 | 37 |
| O | Posttranslation modification, protein turnover, chaperones | 79 | 75 |
| P | Inorganic ion transport and metabolism | 81 | 77 |
| Q | Secondary metabolites biosynthesis, transport and catabolism | 42 | 34 |
| R | General function prediction only | 243 | 202 |
| S | Function unknown | 116 | 101 |
| T | Signal transduction mechanism | 166 | 152 |
| U | Intracellular trafficking, secretion, and vesicular transport | 55 | 43 |
| V | Defense mechanisms | 22 | 20 |
| W | Extracellular structures | 0 | 0 |
| Y | Nuclear structure | 0 | 0 |
| Z | Cytoskeleton | 0 | 0 |

**Table S6**. The putative selenoproteins from the genome that are separated as previously identified and candidate novel ones. Previously identified selenoproteins are shown in bold.

| *#* | *AA length* | *Protein ID* | *Name of protein* |
|---|---|---|---|
| **1** | **69** | **PF02662** | **Methyl-viologen-reducing hydrogenase, delta subunit** |
| **2** | **86** | **PF02662** | **Methyl-viologen-reducing hydrogenase, delta subunit** |
| **3** | **565** | **PF00384** | **Molybdopterin oxidoreductase** |
| **4** | **110** | **PF01206** | **Sulfurtransferase** |
| **5** | **126** | **PF00581** | **Rhodanese-like domain** |
| **6** | **218** | **PF00581** | **Rhodanese-like domain** |
| 7 | 252 | TIGR01083 | Endonuclease III |
| **8** | **1417** | **COG1148** | **Heterodisulfide reductase** |
| **9** | **140** | **PF02662** | **Methyl-viologen-reducing hydrogenase, delta subunit** |
| 10 | 374 | COG0482 | tRNA U34 2-thiouridine synthase MnmA/TrmU |
| **11** | **494** | **COG1148** | **Heterodisulfide reductase** |
| **12** | **582** | **TIGR01591** | **formate dehydrogenase, alpha subunit** |
| **13** | **108** | **COG0374** | **Ni,Fe-hydrogenase I large subunit** |
| **14** | **855** | **TIGR01553** | **formate dehydrogenase-N alpha subunit** |
| **15** | **978** | **COG1148** | **Heterodisulfide reductase** |
| **16** | **78** | **PF02662** | **Methyl-viologen-reducing hydrogenase, delta subunit** |
| **17** | **274** | **PF00581** | **Rhodanese-like domain** |
| 18 | 142 | PF02662 | Methyl-viologen-reducing hydrogenase, delta subunit |
| **19** | **150** | **TIGR03865** | **PQQ-dependent catabolism-associated CXXCW motif protein** |
| **20** | **346** | **TIGR03710** | **2-oxoacid:acceptor oxidoreductase, alpha subunit** |
| **21** | **368** | **TIGR00476** | **selenium donor protein (selD)** |
| 22 | 197 | TIGR02287 | phenylacetic acid degradation protein PaaY |
| 23 | 223 | PF00565 | Nuclease homologue |
| 24 | 186 | PF02283 | Cobinamide kinase / cobinamide phosphate guanyltransferase |
| 25 | 481 | COG2403 | Predicted GTPase |
| 26 | 414 | PF13649 | Methyltransferase domain |
| **27** | **213** | **COG0247** | **Fe-S oxidoreductase** |
| 28 | 706 | COG0642 | Signal transduction histidine kinase |
| 29 | 118 | TIGR00952 | ribosomal protein S15 |
| **30** | **343** | **PF00994** | **Probable molybdopterin binding domain** |
| 31 | 127 | NA | Sulfur reduction protein DsrE / hypothetical protein |
| 32 | 78 | NA | Sulfur reduction protein DsrE / hypothetical protein |
| 33 | 265 | PF17131 | Outer membrane lipoprotein-sorting protein |
| 34 | 367 | PF02195 | ParB-like nuclease domain |

**Table S7** The putative selenoproteins amino acid sequence identified by number. Selenocysteine (U) residue shown in red.

| # | *Amino acid sequence* |
|---|---|
| 1 | MSDTNFEPYIISFLCNWUAYAAADLAGVSRMQYPSNMRVIRVTCSGSVSPHHVLKAF<br>QQGVDGVFVGG |
| 2 | VDGVFVGGUHKGECNYLYGNYSAEKRVTILSQMLDFCGIEKERLRARWVSSVEAPEY<br>IEEINDFVGVLKKLGPSPLKNEQAKVIAx |
| 3 | MQKFARAGVGTNNIDHCARLUHSSTVAGLAASFGSGVMTNSTSELEESDAIFIIGSNTT<br>SSHPLVATRIFRAIEKGAKVLVADPRKNQIADLAHLYVRHKPGTDVALLNGIMKVILD<br>KGLEDKKFIAKSTEDFEVFKKQIDTVSIEKTSQITGVSVEEIQAIAEAYGKAERGSIVYC<br>MGITQHTNGVNNVKSLANLSMLTGNIGRVGTGVNPLRGQNNVQGACDMGGLPNVYT<br>GYQPVTLGPANKKFAEAWGVDELPTNIGLTIPELMHGIEKEEVKALWIMGENPVVSDP<br>DANHVVKALEKIELLIVQDIFLTSTAKLAHVVLPGVSFAEKDGTFVNTERRVSRVRKA<br>VEPVGDSRQDWQIIMDISNRFGFEMAYDSPEAVFNEITELTPSYAGITYERIEGPGIQWP<br>CPSLEHPGTPFLHKDGKFTRGKGLFHAIESKPPAEVVDDEYPFWMTTGRVYAHYHTGT<br>MTRNSKALDNEVPEGFLEIHPEDAKSLDIGDGHKIKMASRRGEIETRITITDRVKKGVV<br>FMPFHFEEAKVNKLTNPAYDPIAKIPELKVCAVKLAKV |
| 4 | MRKIWQAICYTHSHGKLUFENQHLNLRECIVPINRNIDIRDVDWPICILNCTREVNQMK<br>TGEQIEVLVKDIDVVNSIVELIEQLLDHSIEKHKKNKYYRLLIKKKQKDT |
| 5 | MTDKLPKDKNALLIFFCQGPTUKMSHKSAWKAEKLGYTNVKLYAGGFPEWKKAGN<br>YYVVETSYVKNVIDSLKPVVIVDSRPTRPAYVKSHLPGAINIPHKKFDALKGILPTDKNI<br>PLIFYCGGYT |
| 6 | VKNVIDSLKPVVIVDSRPTRPAYVKSHLPGAINIPHKKFDALKGILPTDKNIPLIFYCGG<br>YTUKMSHISARMAGQNGYKNVVVYSAGIKAWTKAVGISSGAPTVKVSGDSEVIDIAV<br>FEKIIKENPESIQLIDVRDKDEYAAGHFKTAVNIPTDDLEKKVKTLSDEKPIVFVCNTGA<br>KSGEAYYMLQDLRSDLKKVYYLDAECKYKKDGTFKIIKPK |
| 7 | MSIPWLUSSRADRILSSPPEQRPRPLTWLAVSSIFIYTHSMILNASKIKTILETLRDRYPV<br>VKTQLEHKTPFQLLIATIMSAQCTDNQVNKVSKNLFKKYPDPEDLGHAPLNDIKRIIYS<br>TGFYNNKAKNIKACALAILNEYQSKVPEDISKLVKLPGVGRKTANVVLSAAFGHQTIV<br>VDTHVLRISRRLGLVKSRDPVKVEYELMQIIPRTSWSDLSLQLIYFGREICDARKPLCED<br>CQLFKLCLIKGKG |

**Table S7 continued.** The putative selenoproteins amino acid sequence identified by number. Selenocysteine (U) residue shown in red.

| # | *Amino acid sequence* |
|---|---|
| 8 | M**U**VISPKLVEVGRHLNIELLTNTQLLELEGEEGNFTARVKEAPRFVDLLKCTSCGECVK<br>VCPVEVPSEHNQGLAPRKAIFKQYEQAIPGAYGISKRSTAPCKATCPAHVSIQGFIALM<br>NQGKHAEALKLFKEEHPFPGSCGRVCHHPCELECTRGDVDEPVAIQYLHRYLAELDFE<br>GDETFIPEVAEKRDEKVAIVGSGPAGLTAAYYMAQKGYGVTIFEKLPIKGGMMAVGIP<br>EYRLPRAELEKEIEVIEKLGVDIRTGVEFGKDITLKSLEKDGYNALFMATGLHGSRGLG<br>VKGEDMDGILKCTNFLRDAALGKAKKLSGKVMVIGGGNVAVDVALTAKRLGADDV<br>TMVCLEKRDEMPAWDYEIEEALEEGVKIVNSLGPLRFLGDNGKANEVEFQECTSVFDE<br>NGRFSPQYDDCRLTSYEADTIIVAIGQMGELDYADKEGIALTPPGGFEADPMTLQTPID<br>WVFAGGDAFYGPKSVVDAVASGKEAAESMHRFINGEDLAKGREKTWDFEKPDILDVP<br>KLERVKPSTISLDEREGNFKEITLALAKEAIDKEAARCLSCGICSECYQCVDVCLAGAIN<br>HEEQEKIRDLNVGSVILTTGATTYDPSGLDDIYMYKRSQNVMTSLEFERILSAGGPTLG<br>HLVRPSDEKEPEKIAWLQCIGSRDTNKCGNGYCSSVCCMYAIKDAMIAKEHSEAELDA<br>AIFNMDMRSFGKDYEKYYNRAADQDGVRFVKSRIHSVVEEPGTDNLILNYADEEGKM<br>HKEIFDMVILSVGLEIPKESVDLARRIDIDLDKYNFVKTQTFNPLQTSRPGVYVSGSFQG<br>PKDIPSSVTEASGSAAAAGMKLAPARHTCTKTVVMPEERDVLGEEPRIGVFICKCGINI<br>AGVIDVEDVENYTKTLPNVVYTGENLFTCSQDTQVSIKELIEEHQLNRVVVASCTPKT<br>HEGIFMDTLEEAGLNKYLFEMANIRNQDSWMHFHEPQKATEKAKDLVRMAVARVAT<br>LGPLHDKRISVIDKALVIGGGIAGMNAAKVLAEQGYEVTLVEKQSVLGGLGNKLHHTI<br>EGDDIRAYVKDLVEAVEHHDKIEILKQALIVGFGGYKGNFETTVLVGPSMEERKIDHG<br>AMIVATGATEYQPTEFLYNESDAVVTQIELTDMIEENKAKDLDRVIMIQCVGSRNEDN<br>PNCSRICCQSAVKNAIALKEQNADTDVFILYRDMRMYSMLEEYYTKARNLGVIFSRFD<br>PENQPEVASSENGKMAVTFTDHVLGQKIEASADLVVLSAGVKAADTEELSTIIKTGRN<br>AEGFFMEAHVKLRPVEAVTEGIFICGTAHGPKLISETITQAMAAASRATTFLSQEYLTLS<br>AVTAEVIQENCASCLVCVRSCPYDVPVMNEMGVSYIDPALCQGCGVCAAECPAKTIKF<br>NWYEDNQLLSKVESLLEGV |
| 9 | VILAFCCNF**U**GYSAADLAGSMRLKIPSNFRIVRVPCTGKVDIIHILRAFEKGADGVYVV<br>GCMEGDCHFNEGNFRARKRVEQAARILDKVGVGGERVKMFNLSSGEGPLFAQYSIEM<br>DNKIKELGPSPIRVAKQNKKDAA |
| 10 | MSS**U**NGVLCSSWVLTTGYLSLNVSRIVFIFDAFNIIEWVYMKIEETASQVKGLGLCSGG<br>LDSILSALLLQSQGIDIIWICFETPFFSSESAKKASCITGIPLITLDITDEYMEMMRNPKAG<br>FGKNMNPCMDCHALMFSKAGEVLEKKGFHFLFSGEVLGQRPKSQNKNSLRYVEKNS<br>GFDGQILRPLCAKLLPETLVEQKGLVDREKLLDISGRSRKIQMQMAQDFEIKEYPAPAG<br>GCLLTDKVFSKRLKDLMNVQKVFDKRELYFLKYGRHFRLDLKTKVIAGRSKDDNKHL<br>LKYFNTDKDILLKHAELPGPDVILTGSLTSKNIQTAAMICAAYTKSKPGETANIKVIKK<br>KKKSIICVKTTSALEFKNLMI |

**Table S7 continued.** The putative selenoproteins amino acid sequence identified by number. Selenocysteine (U) residue shown in red.

| # | *Amino acid sequence* |
|---|---|
| 11 | MUILSPKLVEVGRHPNIEVLTFTEVDTVEGQKGDFQVTLKTRPRYIDEDKCTGCTTCVEYCPVQYPDPFNQDISMNKAVHVYFSQAIPLVSYIDDSCLRLKEKKCDICSSVCKTGAIDFRQVPKKTDINVGAIILSSGITPFDPSAREEYGYTKYQNVVTSMDYERLLSSTGPYEGEVKRTSDKEHPKRIAWIQCVGSRRVTEGDNSYCSGVCCTYTQKQVILTKHHYEDVECTIFHNDIRSFGKDFERFFQRAEQLPGVEFIRSYASIEKEIPETNNVVVRYATADDGVKSQEFDMVVLSVGLNPPKAYKEISEKFGIDLNSHGFAKSESSNPIKTNRPGIFVSGAFQGPTDIPESVFTASGAGAQTGELLNYRRGKLAKERIYPEERNVSNEAPKIGVFVCHCGANISSVVNVPSTVEYALTLPNVVYATEQIFSCATNSAQEITELALEKGFNRVVIAACSPRTLEPLFHDTLREAGLNQYYLDMANIREH |
| 12 | MGVLTSARITNEDNYIAQKFTRAVLKTNNIDHCARLUHSSTVAGLAAAFGSGAMTNTIADIETSDMILVTGSNTTENHPVLSSFVKRAVLKGKKLYVIEPRRIKLTEHATKWLRPTPGTDIAWINGMMHVIIKENLHNKDFIENRTENFEALKETVEKYTPEHVEDITGIKADDLIEVARQFAKAPAASILYCMGITQHTCGTDNVKSLANLSMLCGHLGKPGGGVNPLRGQNNVQGACDMGGLPNVFTAYQLVGNDEARSNYEKIWNTTGMSASPGLPVTEMIQKAYEGDFKSLFVIGENPMVSDPDLNHAKKAIANLDFFVVQDIFQTETTRMADVVLPAVCFAEKNGTFSNTERRVQRVRKAVEAPGETKQDWEIICEIAKRMGYEMSYENSQEIFEEIRSVTPSYKGITWDRIDKIGIHWPCPDETHEGTPILHTAQFTRGKGLFHAIDHTPPAELPDEKYPYILTTGRVLYHYHTGTMTMKTDGLNMLSPECFVEISSNDAQKLGLDTGSMVDVSSRRGKISAKLKVSSKAVDGTIFIPFHFAKAAANELTNAKLDPIAKIPEFKVCAIKIEPGTV |
| 13 | VNVGRLIRSFDPULGCAVHVLDADTGRKIKVEIPL |
| 14 | MRSLGLVYIEHQARIUHSATVAALAESFGRGAMTNHWIDIKNSDCILIMGSNAAENHPISFKYVTQAMEEGAKLISVDPRFTRTSSKADIYASLRSGTDIAFLGGMIKYILDNNLYNKEYTTAYTNASFIVGKKFGFKDGLFSGYKKNEKGPALGSYDKSNWAFEMDADGVPETDRTLKHKRSVLQLMKNHYSRYDLDMVSKVTGTPKEDLLKVYKTYAASGARGKAGTIMYAMGWTQHTVGTQNIRTMAIIQLLLGNMGVAGGGVNALRGESNVQGSTDHCLLWHIWPGYLKTPRSSNNSLDAYNTKWTPKSNDPLSANWWGNYPKYSVSMLKSFFGEAATKTNQFGYNWLPKVDDGKAYSWFDIFDDMYKGDIKGFFAWGMNPACSGSNVSKIRKAMENLDWMVNVNIFDNETGSFWKGPGKDPSKIKTEVFMLPAAVSVEKEGSITNSGRWMQWRYQGPKPLGNSRPDGDIIIELGHRIKEEYKTSGGVFQDPIANLKWDYETDGVYDPHKAAKEINGFFTKDVTVKGKLCKKGTLVPSFAWLQTDGSTSSGNWLYCNSYTEKGNMSARRSKKDAPNNIGLYPEFAWCWPVNRRIIYNRASVDPKGRPWDKKDWVIKFAGDEKDGKYVSKKWVGDVPDGGWYPLENPDGSKRKDAKHPFIMRKHGHAQIFGPGRADGPFSEHYEPLESPVKKNAFGSQRINPTAVVYSTKADAYANNDPKFPIVGTTYRVSEHWQTGLMTRPQEWLMELQPNVFVEMSEELAKLRGIKNGERVNVTSARATLECTAIVTKRFTPFKIDNKTVHQIGIPWHYGWRWPSTGAEESANLLTPPAGDPNTRIPETKAFMVNVIKL |

**Table S7 continued.** The putative selenoproteins amino acid sequence identified by number. Selenocysteine (U) residue shown in red.

| # | *Amino acid sequence* |
|---|---|
| 15 | MUIISPKLVEVGRHINIELLTLSEVQHISGEAGDFKVDIVQHPRFVDMDKCIACGLCA<br>EKCPKKVDDEYDEALGKRKAIYVKYAQAVPLKYSIDATNCIYLTMGKCGKCEEVCP<br>TDAINYKDQPKDITLNVGSIIISSGCKPYDPGEHDVYGYTKSKNIVTSLEFERILSSAGP<br>YEGHLVRPSDKKEPKKIAWLQCIGSRDNHLGSNGYCSSVCCTYAVKEAMLAKEHSH<br>DPLDTAIFYMDIRTHGKDYEHFYNRGKDESGIRFVKSKITNIVPDQKTDTQIIKYIDEA<br>GIRQEEAFDIVVLSVGLCIGDEAVELAGKMNISLDHYNFVTTNSFEPVKTSKPGVFIC<br>GAFEAPKDIPSSVIESSAAAGMAGIDLMDNRWSLTKTKEVPEEINITGEAPRIGVFVC<br>RCGTNIAGVVDVPAVVEMAKKLPYVEFAQENMFSCSQDTQDAITNIIKEKQLNRVVI<br>AACTPKTHEGLFQETLTNAGINKYLFDMANIRNQCSWIHAKEPEKATEKAKDLVRM<br>ITAKVALHEPLKEPTLEINQSGLVIGGGVAGIMAAKTLADQGYHTHLLEKEDKLGGQ<br>ANKLYQTWQGEDIQANLSSMIKSVEENKLIDIHLNTEITDVDGFVGNFETHISENGVS<br>ETLKHGVTIISTGSSELKPEEYLYGEDDRVITGLELDQKFLNNDDDLKSTATAAFIQC<br>VGSRIPDRPYCSKVCCTQSIRNALKLKSINPEMNVFILYRDMRPFGLREDLYTQARTK<br>GIQFIKFDFEKELEVAVNEDQLEILFSDTSLRRKMKIKSDLIVLASSIVPEKKNKLAAL<br>YKVAQNADGFFMEAHAKLKPVDCATDGVFLCGLAHAPKPIDESISQAMAAATRAV<br>TLLAKKKMNMDGTVALVDQEKCSSCGVCVSICPFSAPSFTHEGRYEGKAEINPVLCK<br>GCGLCVASCRSGALHLKGFDNHQIFAQIFALDEIESGDVEQTEPGKEEEKDAATAAV |
| 16 | VSGUHPGECYYLDGNYYARRKFALAGNLLEHMGIEKERLHFSWISSAEATKFVDVV<br>KEVTKIVETVGPNNKLVKIPA |
| 17 | MQFLTPAKSLDFDETNAYISQHPGDKITILDVRQPSEYQESHIPGATLIPLPQLSDRLD<br>ELDPEKPTLVYUAVGGRSRVAAQMLAGKGFKKVFNVAGGIKAWHAKTAIGPQDLG<br>MDLFTGKEEPLDVLKVAYSLEQGLYDFYIVMEKEAEHEKVKDLFGKLSEIEVKHQM<br>AIYTAYNDISDEEPSKDEFEKMVEIKALEGGLSTREYIDLFNPDLTSETDVISLAMSIE<br>AQALDLYQRVSSKIENPQSRDIINQIANEEKAHLASLGKLMDSL |
| 18 | MTQEFEPRIIAFCCQYUAYAAGDLAGSMRLSYSADIKVIQVPCTGRVDILHLLNAIED<br>GADGVYVAGCLEGECHYIEGNLKAKRKVEYVKKTLTELGIEPERVAMYNLSSAQG<br>ARFAEIANEMADKIKALGPTPVKNTHAA |
| 19 | MTYCASGTUWKSYHAARLAVENGYTNVYWMRDGIKTWKEAGYSTEGRLKLLNDL<br>IDVNKINFGALCITEKDAKKLKNCTFVDFRDKSKYEKNHIKGANHVDYSDMFSKPM<br>MEELNKSNSLIIIHDAQTVAGVIATTLKLMDYPDVYILK |
| 20 | VWYLHSLLSQKRTUFIQMEDEIASMGAIIGASLTGNKVMTATSGPGFSLKQEALGYA<br>CMAEIPCVIANVQRGGPSTGNPTHVSQGDVNQARWGTHGDHAIIALTASNHQDVFK<br>ITVDAFNMAETYRTPVIILLDEVIGHMREKLVIPEPGEIPVVERLRTSVKKGIDYHPYL<br>PREDGRLPMSDFGAEHRYNVTGLFHDMWGFPNNNPEVVSELLRHLVDKIENNVHEI<br>SMHKEYWMDDAEYILISYGSSARSAIHLARNRRSRGVKLGVLELQTLWPFPTEMVK<br>EKCAGAKAVIVVEMNMGQVVTQVKNAVDNPQTVFLANRIDGQLISPTDIKSLLRMI<br>QGKGV |

**Table S7 continued.** The putative selenoproteins amino acid sequence identified by number. Selenocysteine (U) residue shown in red.

| # | *Amino acid sequence* |
|---|---|
| 21 | MEKINKVFLTQKVKASGUAAKIDPGTLDRIVGGLKVKSHPNLIVGLDTPDDAGVYKL<br>DSKTALIQTLDFLTPVTDDPYEFGQIAAANSLSDVYAMGGEPVTAMNIVCFPSCDLA<br>EDILAETLKGGLDKINESGATLVGGHSVDDQEFKYGLSVTGIVHPDKVLTNAAAQTG<br>DIVILTKPVGTGVMSTAIKAKLAATQNIKEAFATMSLLNKTAAKVMSKFNVNACTD<br>VTGFGLAGHLLEMAKGSKKCISLYTKKVPLLNNVLDFANMGMVPAGAHKNRNFFN<br>DLTHIDADVNRAVVDLMFDPQTSQSCLLCRWHLPLVLLSCLPCPLQHLWAPWFRLF<br>LKDLILILPLQQDLLSPLPSILPAYIAIL |
| 22 | VQALPPEPFRKLWVIIUWEKGKDMAVYRYGERIPQIGKNSYVSDSARVIGDVTIKDN<br>CYIGHGAIVRGDYGKIIIGHGTAIEENAILHIRPNGILTLEENVTVGHGAIIHGKLIKSQ<br>AVIGIGSIIGFDVVIGSWSIIAEGCVVPQNTIIPDGKITGGVPFKIIGNVKQKHKDFWTY<br>GKQLYVDLAKEYPEKFEKL |
| 23 | MRKNGPGLCWUKKYDNPDRMIKSVVYIFFTLFLSLFLYGNIYFWVDENGIKHFTNIT<br>PALNETVEEFKESNIVFPNQQFKVLKVFDGDTIKVTRVDLIFKIRLVGIDSPELGFKGQ<br>KTQPFSQKAKQHLTGLLKNKKVRIKSYGTDAYNRQLAEVFSGNKNINIEMIRAGLAE<br>VYKGRRPKKLDSQIYLKEEARARKTGKGMWIQGRLYKSPRQWRKEHPRKx |
| 24 | MIUIKHKLLKKNVTFVIGGCRSGKSFFALDQANRIKGGKKYFIATSVPTDTEMEKRVE<br>SHQKERGQDWHTIEEPVMIHEKINQYSTEARVLLVDCLTLWVSNLLFHSYDKTRIDE<br>AVKHLENSLEQCECPIFLVSNEVGCGIVPENELARKFRDFAGFVNQRMAKIADTVVM<br>TIAGIDVQIKPRL |
| 25 | MLKDQPRSGUIDHRSFRCGSILMSLYKTEGIMDKRIRRNVLILGAAGRDFHNFNTFYR<br>DNKNYKVVAFTAAQIPDIDDRKYPVELAGELYPDGIPIYSEEELPRLIKELEVDECEFS<br>YSDVPNQHVMNISSIVNAAGANFVLVGTKDTQVQSNKPLISVCAVRTGCGKSQTSRR<br>IIEHLMAKGLKVVAIRHPMPYGDLVAQKVQRFASLEDIDNQNCTVEEMEEYEPHVTR<br>GNVIYAGADYEAILRAAENDPDGCDVILWDGGNNDFSFYKSDLNVTVVDPHRAGHE<br>LTFYPGTATLRMSDVVVINKMDSADAAGIEQVRKNITALVPDAIVVDGASTLDVEDP<br>GLIKGKKVLIVEDGPTLTHGQMKYGAGTVAAQKFGAAEFVDPRPYTVGRLSETFET<br>YPEIGCLLPAMGYGKQQLKDLETTINNTECDTVVIGTPIDLSRIIKIDKPSTRVIYSLQEI<br>GRPDLGGILADFIDTHNLV |
| 26 | MAKRVGNYFKQKGLSVVTPSNAGHFNYQNTRIYYARGYYEDALELSRKIPGFEIAG<br>KFIESTRLKTDIRVLIGKDIVPFDKELRKIUTRCVGNENSDSKINCEMELIKCFKALSDK<br>TRLRLLYVLQHYELNVNEIVLVVDMIQSGVSRHLKILMESGLLTSRRDGSFIYYSAAK<br>NDAVKTVLSLVDQSLEKEETAGQDLAASREMIKIRQNRTRRFFKTVAPQWDRLKKE<br>VLGDFDLNSMIKEKVCFHGNISDLGCGTGELIEILSEQTSHKLIGIDYSPEMLEQARLR<br>LSGTGNAEIRLGELEHLPMKNQEIDTAVMNMVLHHVSQPELPISEVYRVIKPGGLFIL<br>SDFEKHDKEKIKEIIGGSWLGFEKEKIKTWLTDAGFHLKKIDSYPVNHDLIINVFTAK<br>KPSIQEKL |

**Table S7 continued.** The putative selenoproteins amino acid sequence identified by number. Selenocysteine (U) residue shown in red.

| # | *Amino acids sequence* |
|---|---|
| 27 | VDQANKLGCKTYLNTEUGHVTFSVLAGLKKFKLEHNFDVKNMYEYYAKWIREGK<br>LKVNSDWNKDLKIKFTVQDPCQIVRKSYGDPIAEDLRFVVKSVVGEENFVDMQPNK<br>SNNYCCGGGGGFLQSGFKDQRLAYGKIKDEQIKATGADYCIAGCHNCHAQIHELSE<br>HYGGNYPVVHLWTLICLSLGILGPNEREYLGDDLKEVNVFHPETAM |
| 28 | MITQITUFSKRLWSMTKILAIDDKKDNLVSLSATLKSLIPGCTVITAMSGLEGIEKAKT<br>ESPDTIVLDIKMPGMDGYETCKRLKKNKETGHIPVIMVSAIMTETRGLVKGLDTGAD<br>AYLAKPIDEHVLAAQIKTTLRMKKAEDVLRRQKKVLEDSVLKRTAELTSSNAQLRR<br>EIEERKQTEGSLIKSEEKLSQAIQGNSTPTFIIDNNHTITHWNNACEKLTGLSEATMVA<br>TKKQWLIYYDKERPVLADFIVDGVPGEVIDRYYKRKHQKSILVEGAYEAEQFFPNFG<br>EKGKWIFFTASPLRDHDGNIMGAIETFQNITERKSTEAQLRQAQRMESIGTLAGGIAH<br>DFNNVLFPILGHADMLLADIPEDSPSRDSLNKIYTSALRARDLVRQILTFASQDTNELI<br>LMEMQPVIKEALKLIRSTIPTTIDIKQDIRSDCGTIKADPTQIHQIVMNLATNAYHAME<br>DTGGELKVSLKEVNLRDDEVITPDMTPGVYACLIIADTGTGMDKNVTEKIFDPFFTT<br>KGTGKGTGMGLSVVHGIVNSAGGAIHVYSEPGKSTKFNVYLPVIKSSLEEQKPQVA<br>KLMQGGTERILLVDDEILIAKMLQQILERLGYHVTSLNSSIDALEVFRANSDKFDMVI<br>TDMGMPNMSGDKLSAELIKIRPDIPILICTGFSEKMTEEKIASLNIKGFVLKPVVMKD<br>LAGKIREMLDEN |
| 29 | MEYRIRNHISSDHUQIPCFIYSRQRSFKVVLLAENKEEMIEQFKLHESDTGSPEVQVAI<br>LTHRISYLTEHLKVHTKDHHSRRGLLILVGRRRSLLDYVKKKDVSRYRSLIERLGLR<br>R |
| 30 | MKUMKKIRVENAVGTVLAHDMTRIIRGKFKGVGFKKGHVIQKKDVPELLKIGKQY<br>LYVLDLGKNQLHEDDAAKRIAKAISDRDLDFSEPREGKINISTPYAGLLKINVDALLQ<br>VNKMESIIVATLKNNFPCKKGEIIAGTRIIPLTIPAKNIEALESLAKETGTILWVKPYKS<br>LKIGAVVTGSEVFNGLITDDFGPSVGKKITDAGCTLIKKIIVPDEEKAISNAILELKQL<br>GCEMIITTGGLSVDPDDVTRQGVIRSGANVTFYGSPVLPGAMLMVSHLADIPIISLPA<br>CVFYYKQTVFDLIFPRVLAGETISEDDIAAMGHGGLCMNCKVCHYPVCSFGR |
| 31 | MVFAURENSGCPSQWRDLYSSERYQSDTNLYQPNKKYSYKRRETPNQPGLKFFKGL<br>FMKIAIVICQDIPEVLWNAFRLANMMLEGMEDVTIFLNGPSVKYEELDSTQFPLIELS<br>KIFTLSEGELLA |

**Table S7 continued.** The putative selenoproteins amino acid sequence identified by number. Selenocysteine (U) residue shown in red.

| # | *Amino acids sequence* |
|---|---|
| 32 | MEDVTIFLNGPSVKYEELDSTQFPLIELSKIFTLSEGELLAUGKLIDLHGVKEGYHKSG<br>TQKNLYDLIKESDKIITF |
| 33 | MKVDSICIUGGKMRKAMLIVVILFFAVSSQAYAQDRTNAETIVKKAFDYWRGDASV<br>STITMTIHRPDWERSMTIKAWTRGESDSLFVITNPAKDRGNSTLKAGKGMWMYNPK<br>VNRVIKLPPSMMSQGWQGSDFSNNDLAKTDSLIKDYVHTLEDTKVDDGKKVYSIKS<br>MPKPDAPVIWGMIKLTIREDNILLREEFFDEDFMSVKIMTAWDIQMTGSKLFPMKWK<br>MQKSDADNEYTQFVYEKIDFIKSLSKNIFTRTYLKNPSI |
| 34 | MSGTGSLIPDGIKNIRAPNCPGKLLIRQQRHIKTFSHVUQVKMSDLLCKINILDIDLTD<br>KRYKISFAEDDITFLAHSIKETGLITPPVVRPLNNKFVIISGFNRIRALIYNNRLYNNET<br>KIVVYKTKPDITDCSCLVKAIAALAFKRPLTHSELIISTRRLYQFLDKKQIAKKSAAIF<br>NIEFNVRFVEDLLTIGALPDPAFELIHSGNLSFKSAKRISSYGENIIKVFLTIFSKIKASS<br>NNQLEIILHIMEISARDAIKPESLLKNKEMQSILFDENKDPGLKTKDLRAWLFEQRFPT<br>IFKAHQMVREKITSIKFKNNIKFLPPQNFESQNFSISFTAKNYSEFAASVQNLNTALEN<br>RELKEIFNQ |

**Table S8**. Genes proximal to trimethylamine methyltransferase in Deltaproteobacterial-bbl genomes.

| TIGRFAM | Deltaproteobacteria -bbl-31 contig no. | Protein |
|---|---|---|
| TIGR02369 | c_000000000882_1 | trimethyl_pyl: trimethylamine:corrinoid methyltransferase |
| TIGR01096 | c_000000000882_2 | lysine-arginine-ornithine-binding periplasmic protein |
| TIGR03004 | c_000000000882_3 | ectoine_ehuC:ectoine/hydroxyectoine ABCtransporter permease protein EhuC |
| TIGR03005 | c_000000000882_4 | ectoine_ehuA: ectoine/hydroxyectoine transporter ATP-binding protein EhuA |
| TIGR03338 | c_000000000882_5 | phnR_burk: phosphonate utilization associated transcriptional regulator |
| NA | c_000000000882_6 | glycine betaine methyltransferase |

**Table S9**. Locations, accession numbers, and other information from the data sets that were used for this study.

| *#* | *Location* | *Accession* | *Submission* | *Data type* |
|---|---|---|---|---|
| 1 | San Pedro Basin, California | *TBD* | University of California, Santa Barbara | metagenomic / genomic |
| 2 | Landsort Deep, Baltic Sea | PRJEB6616 | European Nucleotide Archive (ENA) | metatranscriptomic |
| 3 | Columbia River Estuary, Washington, USA | PRJNA441934 | DOE Joint Genome Institute | metatranscriptomic |
| 4 | Hydrothermal Vent, Guaymas Basin | PRJNA362212 | The University of Texas at Austin | metagenomic / genomic |
| 5 | Mud flat, Arcachon, South-west France | PRJNA182447 | DOE Joint Genome Institute | metagenomic / genomic |
| 6 | Petroleum reservoir, North Sea | PRJEB18182 | DOE Joint Genome Institute | metagenomic / genomic |
| 7 | Marsh, Singapore | PRJEB16283 | DOE Joint Genome Institute | metagenomic / genomic |
| 8 | Brackish spring, Death Valley, CA, USA | PRJEB14757 | CNRS France | metagenomic / genomic |
| 9 | Marine Mud, Venice, Italy | PRJEB20333 | DOE Joint Genome Institute | metagenomic / genomic |
| 10 | Gutless marine worm, sediment, Italy | PRJNA17779 | DOE Joint Genome Institute | metagenomic / genomic |
| 11 | Petroleum seeps, Eastern Gulf of Mexico | PRJNA485648 | University of Calgary | metagenomic / genomic |
| 12 | Coal Oil Point seeps, Santa Barbara, CA, USA | PRJNA366139 | DOE Joint Genome Institute | metagenomic / genomic |

**Table S10.** TIGRFAM identity and gene names of proteins relevant for genetic code expansion.

| TIGRFAM ID | GENE ID |
|---|---|
| TIGR03912 | PylS_Nterm: pyrrolysine--tRNA ligase, N-terminal region |
| TIGR02367 | PylS_Cterm: pyrrolysine--tRNA ligase, C-terminal region |
| TIGR03910 | pyrrolys_PylB: pyrrolysine biosynthesis radical SAM protein PylB |
| TIGR03909 | pyrrolys_PylC: pyrrolysine biosynthesis protein PylC |
| TIGR03911 | pyrrolys_PylD: pyrrolysine biosynthesis protein PylD |
| TIGR00475 | selB: selenocysteine-specific translation elongation factor |
| TIGR00474 | selA: L-seryl-tRNA(Sec) selenium transferase |
| TIGR00476 | selD: selenide, water dikinase |
